# Fluid transport properties dominate blastocyst expansion over mechanics

**DOI:** 10.64898/2026.10.05.756743

**Authors:** Louise Dagher, Matteo Bassanini, Ludmilla de Plater, Giacomo Gropplero, Clément Caporal, Aurélien Maillot, Ozan Kastas, Charlie Duclut, Stéphanie Descroix, Jean-Léon Maître

## Abstract

Lumens populate most of our organs and their size, which dynamically changes throughout the day and lifetime, can influence their function. During mammalian development, the first lumen appears in the blastocyst. Surface epithelial cells pump ions to build an osmotic gradient that draws water, which inflates the embryo and stretches the epithelium as well as the zona pellucida, an elastic shell protecting the embryo. Blastocyst expansion is not steady and instead undergoes cycles of expansion interrupted by collapses during which the lumen leaks out. Early studies identified the critical role of osmotic pressure difference in regulating the expansion of the blastocyst whereas recent studies proposed that mechanical stresses would set the size of the embryo. However, the relative contributions of osmotic and mechanical stresses to lumen expansion dynamics remain unclear. Here, we investigate this using microfluidics, biophysics, machine learning and modelling on mouse embryos. Breaking down the expansion dynamics of mouse embryos, we discovered that fluid pumping rate accelerates, compensating for the increased frequency of collapses interrupting episodes of expansion. We reasoned that fluid transport could be rate limited by osmotic pumping, water permeability and/or mechanical constraints. Removing the mechanical constraint from the zona pellucida only marginally affected these expansion dynamics. Forcing fluid exchange with osmotic shocks we measured that water permeability is large enough not to restrict fluid accumulation rates. Finally, we estimate ion pumping for mouse blastocysts and interfere with it using inhibitors to measure how it quantitatively sets blastocyst expansion rates. T£ogether, we propose that osmotic pressure differences, regulated by fluid transport properties, dominate over mechanical stresses in controlling blastocyst expansion dynamics.

## Introduction

From brain ventricles to salivary glands, most organs of the mammalian body contain fluid-filled cavities called lumens that are critical to organismal structure and function (Dagher et al., 2024; Lee et al., 2026). Lumens occupy a significant portion of the volume of organs and thereby contribute to setting their shape and size (Chan and Hiiragi, 2020; Yang et al., 2021; Lee et al., 2026). Abnormal lumen growth can result in the formation of benign or pathological cysts that can grow up to tens or even hundreds of liters (Grubor et al., 2013; Shah et al., 2014). Lumens can also determine the size of whole organisms, such as cnidarians, and control their development and behavior (Stokkermans et al., 2022; Weevers et al., 2025). This is also the case for the first lumen of mammalian embryos, called the blastocoel, forming during preimplantation development.

In mouse embryos, the blastocoel forms three days after fertilization when the embryo reaches the 32-cell stage (Płusa and Piliszek, 2020; Rozema and Maître, 2024). At that stage, surface cells have differentiated into trophectoderm (TE), an epithelial extraembryonic tissue that is apico-basally polarized and sealed with tight junctions (Korotkevich et al., 2017; Zenker et al., 2018). Polarized transport of osmolytes (primarily sodium ions) into the intercellular space coupled to the sealing of tight junctions (reducing their leakage) builds an osmotic gradient that draws water from the environment into the embryo (Biggers et al., 1978; Powers et al., 1977). As fluid accumulates between cell-cell contacts, pressurized pockets form and exchange fluid at a distance until all fluid collects into a single large lumen (Dumortier et al., 2019; Schliffka et al., 2024). This lumen then continues growing until the embryo comes into contact with the zona pellucida (ZP), the eggshell from which the embryo will hatch before implantation (Leonavicius et al., 2018). After reaching the ZP, the embryo will continue to grow rapidly thanks to the expansion of the lumen. Lumen expansion is not continuous but instead alternates between longer phases of lumen swelling and shorter episodes of collapse driven by the transient leakage of the lumen content through temporary holes in the TE associated with cytokinesis (Leonavicius et al., 2018; Chan et al., 2019; Guo et al., 2025). Intermittent lumen inflation is actually found in other biological contexts such as the zebrafish inner ear (Swinburne et al., 2018), cnidarian polyps (Fütterer et al., 2003; Stokkermans et al., 2022; Weevers et al., 2025) or cystic organoids (Ruiz-Herrero et al., 2017). Together, this sequence of events highlights the intricate dynamics of molecular, cellular and physical changes leading to the formation and expansion of the first mammalian lumen (Chan and Hiiragi, 2020; Dagher et al., 2024). Understanding what controls the expansion and size of the blastocoel is critical since several studies provided evidences it influences lineage specification (Chan et al., 2019; Ryan et al., 2019; Guo et al., 2025) or hatching after thawing (Leonavicius et al., 2018), which is important to Assisted Reproductive Technologies (ART, Firmin and Maître, 2021).

Early studies on the regulation of blastocoel expansion had emphasized the role of osmotic transport in regulating blastocoel size (Borland et al., 1977; Manejwala et al., 1989; Pedersen and Spindle, 1980; Powers et al., 1977; Rose and Watson, 2007). In fact, osmotic shocks of bovine embryos carried out at room temperature reported that blastocysts behaved as perfect osmometers (Mazur and Schneider, 1986). In that case, the surface epithelium behaves as an ideal semi-permeable membrane, allowing the passage of water to equilibrate the osmolarity between the two compartments it separates by adjusting their volumes. When volume changes scale linearly with osmolarity, the material is said to behave as a perfect osmometer as this follows the van’t Hoff equation that resembles the law of perfect gas (van’t Hoff, 1884; Olver and Benson, 2023). Epithelia being complex materials, their ability to stretch linearly to accommodate inflating lumens appears unlikely and studies reported that epithelial stretching can result in exotic material properties and cell behaviors (Latorre et al., 2018; Golde et al., 2026). Instead, recent studies proposed that blastocoel expansion and size are primarily regulated by mechanical properties (Ruiz-Herrero et al., 2017; Leonavicius et al., 2018; Chan et al., 2019; Chan and Hiiragi, 2020). Indeed, the blastocyst would inflate until it reaches a threshold tension causing epithelial junctions to fail and the embryo to collapse (Chan et al., 2019). Mechanical reinforcement of epithelial junctions would then allow embryos to reach higher tension values and a larger steady-state size (Chan et al., 2019). In this model, osmotic pumping of fluid is implicitly taken into account and is at the origin of lumen inflation between collapses (Ruiz-Herrero et al., 2017). However, increased osmotic pumping has not been considered as an alternative or complementary mechanism of embryo size control (Dagher et al., 2024). Therefore, while both transport and mechanical properties of developing blastocysts have been proposed to set the size of the first mammalian lumen, their relative contributions have not been characterized.

Using time-lapse microscopy and machine learning-based segmentation, we quantify the expansion dynamics of mouse preimplantation embryos during lumen formation. This reveals that fluid pumping accelerates during lumen formation despite frequent collapses temporarily halting lumen expansion. To understand the origin of this surge, we characterize the transport properties of mouse embryos using a dedicated microfluidic device to perform high spatio-temporal resolution imaging of embryos during rapid osmotic shocks. This shows that, despite their complexity, mouse embryos can be approximated to ideal semipermeable membranes where lumen expansion is dominated by osmotic pressure difference while mechanical contribution is negligible. Furthermore, we uncover that fluid accumulation in the lumen scales linearly with time, which we attribute to a linear increase of the lumen osmolarity. Using theoretical modelling of these experiments, we can extract key epithelial transport properties such as osmotic pressure difference, water and osmolyte permeabilities, and rate of active pumping. To test our conclusions, we remove the mechanical influence of the ZP, which does not change lumen growth dynamics or transport properties, and we block ion transport, which quantitatively affects the osmotic pressure built by embryos and their ability to inflate their lumen. Together, this interdisciplinary approach reveals the simple physical mechanisms controlling mouse blastocyst expansion, which is dominated by epithelial fluid transport properties rather than mechanics.

## Methods

### Preimplantation embryos handling

#### Handling of mice and isolation of embryos

All mouse work is performed under the supervision and permission of the institutional veterinarian and in agreement with governmental approval (APAFIS numbers 39490-2022111819233999 v2). The animal facilities are operated according to international animal welfare rules.

Superovulation of female mice was induced by intraperitoneal injection of five international units of pregnant mare’s serum gonadotropin (Ceva, Syncropart), followed by intraperitoneal injection of five international units of human chorionic gonadotropin (MSD Animal Health, Chorulon) 44–48 h later.

Embryos were recovered between E0.5 and E3.5 by dissection or flushing of oviducts and at early blastocyst stage by flushing oviducts and uteruses with 37°C homemade KSOM HEPES (Biggers et al., 2000; NaCl (Honeywell Research Chemicals, Fluka Cat31434), KCl (Sigma, BioReagent P540 LifeGlobal, ZEKS-050), KH_2_PO_4_ (Avantor Science Central, 26936.293), Na Lactate (Sigma, L4263), Na pyruvate (Sigma P4562), CaCl_2_*⋅* 2H_2_O (FisherScientific, C79-500), MgSO_4_*⋅* 7H_2_O (Sigma, 230391), Na_4_EDTA*⋅* 4H_2_O (Sigma, E-5391) MEM essential AA solution (FisherScientific GIBCO, #11130-036), non-essential AA solution (FisherScientific GIBCO, #11140-035), L-Glutamine (FisherScientific GIBCO, #25030-032), D(+) glucose (Sigma, G-6152), NaHCO_3_ (Sigma, S-5761), Penisilin and Streptomycin (FisherScientific GIBCO, 15140122), 4-(2-hydroxyethyl)-1-piperazineethanesulfonic acid (HEPES, Sigma, H4034)) using a modified syringe (Acufirm, Berlin, Germany, 1400 LL 23).

To remove surrounding cumulus cells from embryos collected at E0.5, they were treated for 10 s with 37°C 0.3 mg/mL hyaluronidase (Sigma, H4272-30MG). Embryos are then washed again with KSOM-HEPES and cultured in KSOM (homemade) containing 1 mg/mL BSA (Sigma Aldrich M8410) in 10 μL droplets covered in mineral oil (Sigma, M8410 or Acros Organics) at 37°C and 5% CO_2_ in a humidified atmosphere.

Embryos are handled using an aspirator tube (Sigma, A5177-5EA) equipped with a glass pipette pulled from glass micropipettes (Blaubrand intraMark or Warner Instruments).

For experiments with embryos without zona pellucida, E2.5 embryos are incubated in 5 mg/mL pronase (Sigma Aldrich P8811) for 35 s at 37°C, then washed and incubated in KSOM-HEPES until complete digestion of the zona pellucida.

For imaging, embryos are placed in 5 cm glass-bottom dishes (MatTek).

#### Mouse lines

Mice are used from 5 weeks old on. (C57BL/6xC3H) F1 hybrid strain is used as wild-type. To visualize plasma membranes (Gt(ROSA)26Sor^tm4(ACTB-tdTomato,-EGFP)Luo)^ mice (mTmG; RRID IMSR_JAX:007676) are used (Muzumdar et al., 2007).

### Microfluidic chips

#### Design

To probe the dynamic response of embryos to osmotic shock, we designed a dedicated microfluidic device that meets several requirements: (i) single-embryo hydrodynamic trapping with limited mechanical constraints while maintaining viability, (ii) rapid medium exchange, (iii) compatibility with high-temporal resolution imaging. To fulfil these requirements, the microfluidic device comprises several functional components (Fig S1A). Three symmetrically arranged inlets were designed to first load and trap an embryo and then to inject two media of different osmolarity. The dimensions of the embryo trap were determined through numerical simulations while accounting for microfabrication constraints. The resulting design consists of a trap measuring 150 µm in width and height, connected to a 50 µm-wide channel, enabling efficient embryo trapping while accommodating the volume changes associated with hyper-osmotic and hypo-osmotic shocks. When the trap is empty, the chip geometry ensures that hydrodynamic forces at the constriction guide the embryo into the trap (Fig S1B). Once the embryo is trapped, the flow streamlines are redirected through the 30 µm-wide side channels. In such experimental conditions the Reynolds number is low, the flow is laminar, so the two media flow side by side, with the position of their interface controlled by the pressure difference between the two inlets (Fig S1C-D). As a result, the medium surrounding the trapped embryo can be exchanged without replacing the entire dead volume of the microfluidic chip, substantially reducing the switching time of medium. In this configuration, an overall pressure difference of about 20 mbar is sufficient to exchange the medium surrounding the embryo in less than 30 s, while limiting the mechanical stress applied on it.

We used the COMSOL Multiphysics 6.2 software (COMSOL AB, Stockholm, Sweden) to model fluid flow within the microfluidic device. The simulations were conducted using a 2D model with the default fine physics-controlled mesh provided by COMSOL.

#### Fabrication

Molds obtained by photolithography were first silanized and then treated with oxygen plasma (Diener Electronic PICO-PCCE, settings: 30 s, 0.40 mbar, 100%) and exposed to vapor phase silanization using Trichloro(1H,1H,2H,2H-perfluorooctyl)silane (Sigma-Aldrich, 448931). Molds were extensively washed with 70% ethanol prior to use to prevent cytotoxic effects of residual silane. Chips were fabricated by casting polydimethylsiloxane (PDMS, Sylgard 184, Farnell, 101697) on the mold at a 10:1 elastomer to curing agent weight ratio. Bubbles were removed with a desiccator before curing the PDMS at 75°C for at least 3 h. After peeling from the mold, inlets and outlets were created with a 0.75 mm hole puncher (World precision instrument, 15453532). Devices were cleaned with 70% ethanol and dried with pressurized air. Any remaining particle was removed from the PDMS surface using transparent adhesive tape (Scotch Tape, 3M). Chips were bound to 0.17 mm or 1.7 mm thick cover glasses after treatment with oxygen plasma (Diener Electronic PICO-PCCE, settings: 30 s, 0.40 mbar, 100 %).

#### Embryo mounting and recovery

The chip is rinsed with 70% ethanol, followed by a solution of polydimethylacrylamide-co-allyl glycidyl ether (PDMA-AGE at 0.5 w/v was provided by M. Chiara group from the Instituto of Science and Technology (SCITEC-CNR, Milano, Italy)) for 15 min. The chip is first rinsed with water, then filled with KSOM-HEPES. The medium is then incubated at 37°C and 5% CO_2_ in the microscope chamber. Medium inlets are connected to a bubble trap (Fluigent, CTQ-006BT), and to a M-SWITCH Microfluidic bidirectional valve (Fluigent, ESSMSW003) that relays to independent medium reservoirs. The embryo is carefully placed into the embryo inlet of the chip. The embryo is then injected using negative hydrostatic pressure. When the embryo is captured within the hydrodynamic trap, KSOM HEPES with 10 kDa fluorescently labelled AlexaFluor-647-Dextran (Invitrogen, D22914) is introduced at 15 mbar with a positive pressure controller (Fluigent, Switch EZ). The embryo inlet is then sealed with tape. For dynamic medium changes, both medium inlets are used with KSOM HEPES containing a 10 kDa fluorescently labelled Dextran: either AlexaFluor-647 (Invitrogen, D22914) or AlexaFluor-488 (Invitrogen, D22910) at 1:1000. Fast medium change is obtained with a configuration where the two media flow in parallel in the chip but only one surrounds the embryo by careful positioning of the interface between the two media. Their interface position is controlled by the ratio of inlet pressures (typically ΔP_1_ = 6 mbar; ΔP_2_ = 12 mbar). The interface is swiftly moved by changing pressures (ΔP_1_ → ΔP_2_ and ΔP_2_ → ΔP_1_ ). After experiments, pressures are reversed to retrieve the embryos, which are placed back in a culture dish for observation.

### Perturbations

#### Osmotic shocks

For hyperosmotic shocks, we used solutions containing 43.25, 87.5, and 175 mM sucrose diluted in KSOM HEPES supplemented with 10 kDa fluorescently labelled Dextrans (either AlexaFluor-647 (Invitrogen, D22914) or AlexaFluor-488 (Invitrogen, D22910) at 1:1000) with final osmolarities of 295, 345, and 445 mM, respectively. For hypo-osmotic shocks: 10%, 30% and 50% dilutions of KSOM HEPES supplemented with 10 kDa fluorescently labelled Dextran (either AlexaFluor-647 (Invitrogen, D22914) or AlexaFluor-488 (Invitrogen, D22910) at 1:1000) with ultra-pure water (Milli Q EQ 7000), with osmolarities of 234, 169 and 123 mM, respectively. Osmolarities were verified using an osmometer (Loeser, TYP15).

#### Drug treatments

5-(N-Ethyl-N-isopropyl)-amiloride (EIPA, Sigma, A3085) 50 mM DMSO stock is diluted to 7.5 or 15 µM in KSOM. Ouabain (Tocris, 1076 ) 100 mM DMSO stock is diluted to 150 or 500 µM in KSOM. A 3:2000 or 1:200 DMSO in KSOM dilution is used as control. E3.5 blastocysts with a volume greater than 300 pL were transferred to medium containing either drug or DMSO.

### Microscopy

Live imaging of embryo development is performed using a Cell Discoverer 7 (Zeiss) equipped with PLAN-APOCHROMAT 20x/0.95 autocorr objective, a magnification lens for 2x and an ORCA-Flash 4.0 camera (C11440, Hamamatsu). The microscope is equipped with an incubation chamber to keep the sample at 37°C and supply the atmosphere with 5% CO_2_.

To follow lumen growth dynamics, embryos are monitored from E3.0 (before blastocoel formation to fully expanded state) for ∼15-70 h using bright field imaging in a time-lapse acquiring every 20 min 6 focal planes spaced 12 µm apart at multiple positions tiled next to each other. Tiles are then stitched together in Zen Blue (Zeiss).

Imaging of osmotic shocks is performed on an inverted Zeiss Observer Z1 microscope equipped with a CSU-X1 spinning disc unit (Yokogawa). Embryos are imaged using 405, 488, 561, and 642 nm laser lines through a 40x/0.95 Plan Apo Korr dry objective; emission is collected through 450/50 nm, 525/50 nm, 595/50 band pass or 610 nm low pass filters onto an ORCA-Flash 4.0 camera (C11440, Hamamatsu). The microscope is equipped with an incubation chamber to keep the sample at 37°C and supply the atmosphere with 5% CO_2_.

Embryos in culture dishes are imaged before and 10 min after osmotic shocks with a z-stack consisting of 100 focal planes spaced 1 µm apart.

Embryos in the microfluidic chip are imaged every 15 s with z-stacks consisting of 6 focal planes spaced 12 µm apart.

### Image and data analysis

#### Design and implementation of a segmentation model for bright-field microscopy

First, to generate annotations from bright-field images, the manual drawing tool from Napari (Sofroniew et al., 2026) is used. Three labels are created for each image: embryo, zona pellucida and background. Then, to segment the embryo and its zona pellucida, a U-Net model was trained using yapic 1.2.6 [https://github.com/yapic/yapic] on 1 gpu: Tesla V100 workstation.

At the first image of each time series, the sample is manually annotated, filled to eliminate internal holes and tracked by comparing the position of centroids. For each timepoint, the embryo and zona pellucida automatic segmentation are proofread to eliminate holes or filter out erroneous objects. The area of the segmented embryo is used to calculate its volume assuming it is spherical.

From the evolution of embryonic volume over time inflation and collapse episodes are identified using the MATLAB *findpeaks* function ignoring peaks smaller than 25 pL and considering a 30 min time window during which only one peak can occur. We calculate the slopes, amplitude changes and elapsed time between individual inflation and collapse episodes. Then, we align asynchronous embryos to the time when the embryo reaches 300 µL for the first time or to the time addition of inhibitors.

#### Design and implementation of a segmentation model for measuring luminal and embryonic volumes

To quantify luminal and embryonic volumes, the workflow consists of image preprocessing, deep-learning-based segmentation using nnU-Net, and post-processing of the predicted masks [https://gitlab.curie.fr/pict_bdd/maitre_louise_3d_cells_and_lumen_segmentation/-/tree/main/].

First, a FIJI (Schindelin et al., 2012) macro is used to automatically detect the embryo, crop a standardized 1000 x 1000 pixels region centered on the embryo, and down sample the images fourfold in the x and y dimensions while preserving the original sampling along z. These standardized images are used to train a nnU-Net model for 3D segmentation using biom3d v0.0.33 [https://github.com/GuillaumeMougeot/biom3d/releases ]. Three-dimensional manual ground-truth segmentations of the lumen and embryonic cells are generated for 10 embryos using Napari. Five embryos are used for training, two for validation, and three for independent testing. The model is trained for 500 epochs with a batch size of 4 and a patch size of 80 × 160 × 160 voxels. Data augmentation is enabled, and training uses a combined Dice and binary cross-entropy loss function (Dice-BCE). Segmentation performance is evaluated using the Dice similarity coefficient by comparing the predicted masks with the corresponding manual ground-truth segmentations. The mean Dice scores were 0.935, 0.894, and 0.889 for the training, validation, and independent test sets, respectively. A FIJI macro is subsequently used to retain the largest 3D lumen and cell components and upsample the masks fourfold in x and y.

Using Matlab, the segmented embryos and lumens were closed and filled before extracting the total number of pixels for each 3D structure and calculating the actual volumes. For each embryo, the ratio of final to initial volumes after osmotic shock is calculated. The Matlab function *corrcoef* was used to evaluate the linear relationship between volume ratios and osmotic shock intensity.

#### Analysis of embryo imaging specific to microfluidic chip-based experiments

To detect changes in fluorescent media during osmotic shocks on the microfluidic chip, we use Matlab to measure the total intensity of each fluorescent channel over time and calculate their derivative.

To estimate embryo volume, we segmented maximum intensity projection of mTmG embryos, by applying a Gaussian filter, sharpening images with unsharpen function and using an active contours technique.

An ellipse fitting is performed on the mask to measure the embryo area over time and calculate its volume assuming it is spherical.

#### Statistical analysis

Statistical tests were performed using Matlab and Prism. When comparing different conditions, data distribution was systematically tested for normality and variances were compared. The appropriate statistical test was then used. Differences were considered statistically significant when *P < 0*.*05*.

Sample sizes were not pre-determined and simply result from the repetition of experiments. No samples were excluded. No randomization method was used. The investigators were not blinded to the conditions of the experiments during data collection and analysis.

#### Code availability

Codes for image processing and analysis is available at the following link: https://github.com/MechaBlasto/Dagher

## Results

### Distinct phases of lumen expansion dynamics

To quantitatively analyze lumen growth dynamics, we imaged mouse preimplantation embryos between E3.0 and E4.5, before lumen opening and beyond hatching, and developed a machine learning-based classifier to segment the embryo, ZP and outside medium (Fig 1A, Movie 1). This allowed us to estimate the embryo volume and ZP thickness over time (Fig 1B). As previously measured (Wassarman and Mortillo, 1991; Yu Sun et al., 2003), before lumen opening, embryos with a volume of about 200 pL do not contact the ZP, which is about 9 µm thick (Fig 1B). Lumen formation is initially associated with the appearance of multiple small lumens that coarsen into a single large lumen (Dumortier et al., 2019; Schliffka et al., 2024) and, by the time embryos reach about 300 pL in volume, they eventually come into contact with the ZP, which starts to thin (Fig 1B).

**Figure 1:**
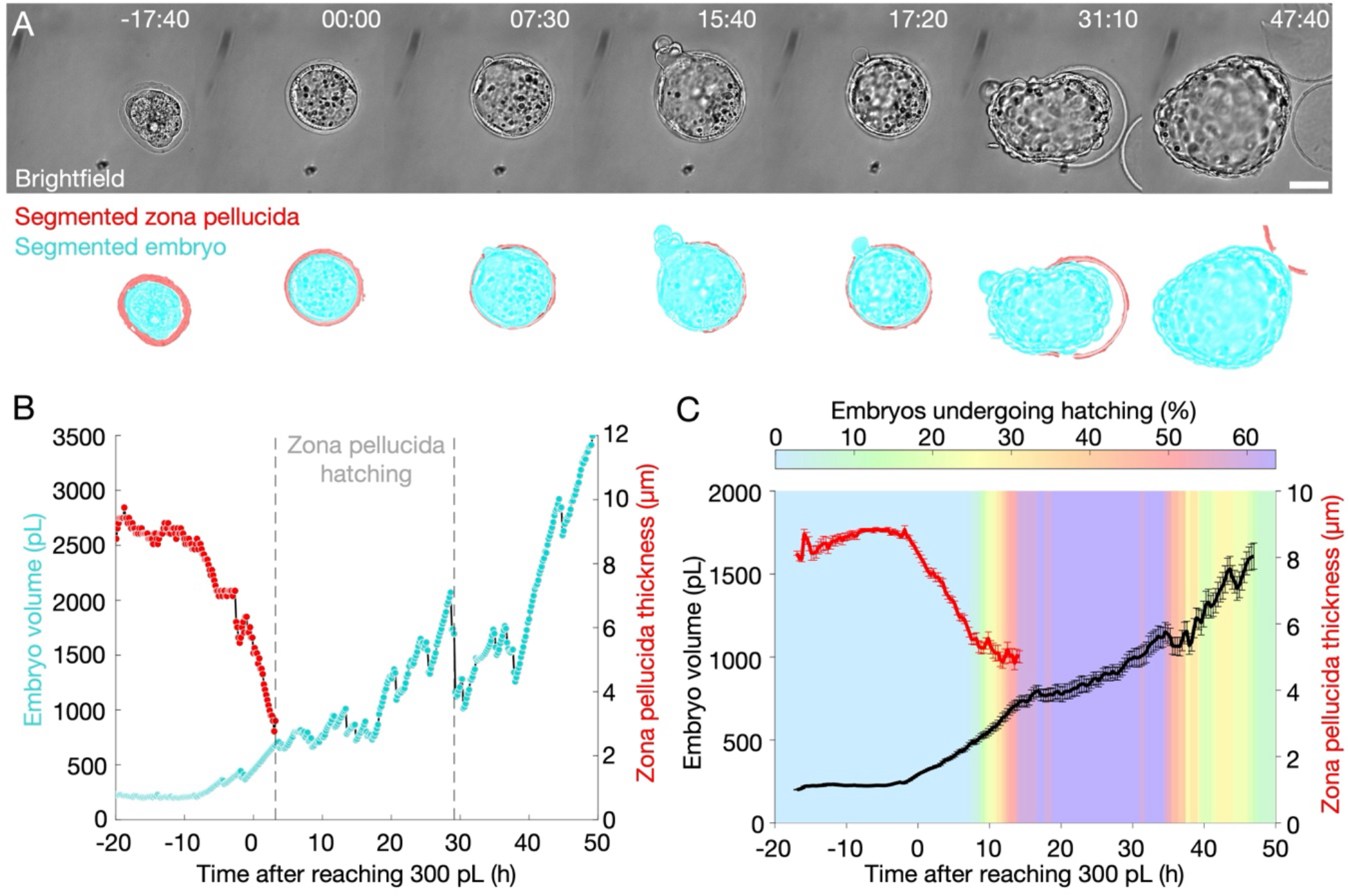
Analysis of blastocyst expansion dynamics reveals distinct phases. **(A)** Top: Brightfield images from a time-lapse of a mouse embryo from E2.5 onwards (time hh:mm, scale bar, 50 µ m). Bottom: Segmentation masks for the embryo (cyan) and its zona pellucida (ZP, red) overlaid on brightfield signal (grey). **(B)** Inferred volume (cyan) and ZP thickness (red) of the embryo shown in (A). **(C)** Averaged embryo volume (black) and ZP thickness (red) of 26 embryos from 3 replicates aligned to the time they reached 300 pL (Mean ± SEM). Background color shows the percentage of embryos that are hatching.

To be able to compare embryos, which vary in their developmental timings, we synchronized them to the moment they reach 300 pL in volume, a landmark corresponding to the time of definite coarsening into a single lumen and of initial contact of the embryo with the ZP. This analysis reveals an initially fast expansion up to about 680 ± 30 pL for about 12 h, before shifting to a slower inflation over the following 36 h (mean ± SEM, 26 embryos, Fig 1C). Interestingly, these phases coincide with the initiation of hatching (Fig 1C, Movie 1) with the majority of embryos initiating hatching about 12 h after reaching 300 pL. By the end of our analysis, most embryos have hatched (21/26 embryos) and their volume exceeds 1500 pL on average (Fig 1C). Importantly, while by that time, *in vivo*, embryos would have implanted, *in vitro* embryos do not show any signs that expansion would stop to reach a steady-state size.

Together, this quantitative analysis of mean blastocyst expansion reveals an initially fast expansion before transitioning to a slower expansion when embryos begin to hatch from their ZP. While slower, this expansion does not stop and by the end of our analysis embryos have expanded from 200 to 1500 pL. With cellular volume now composing a minor part, the blastocyst contains mostly luminal fluid.

### Fluid pumping accelerates and ensures lumen expansion despite frequent collapses

To understand what controls each phase of lumen expansion dynamics, we analyzed the distinct episodes occurring during the formation of individual blastocysts: uninterrupted expansion and collapses (Fig 2A). From the dynamic changes of individual embryo size over time (Fig 1B), we identified automatically the events of major transient leaks leading to collapse and the timing of subsequent re-expansion. With this information, we could determine the collapse frequency and duration, as well as the leak flow and net loss of volume per collapse (Fig 2A). We measured that the frequency of collapses is initially low, with about one every 5 h (Fig 2B). Then, 10 h after reaching 300 pL, this frequency accelerates sharply to about one collapse every 2 h. Therefore, the initially fast expansion measured after averaging embryos may be explained by the lack of interruption by sudden leaks and embryo collapse (Fig 1C).

**Figure 2:**
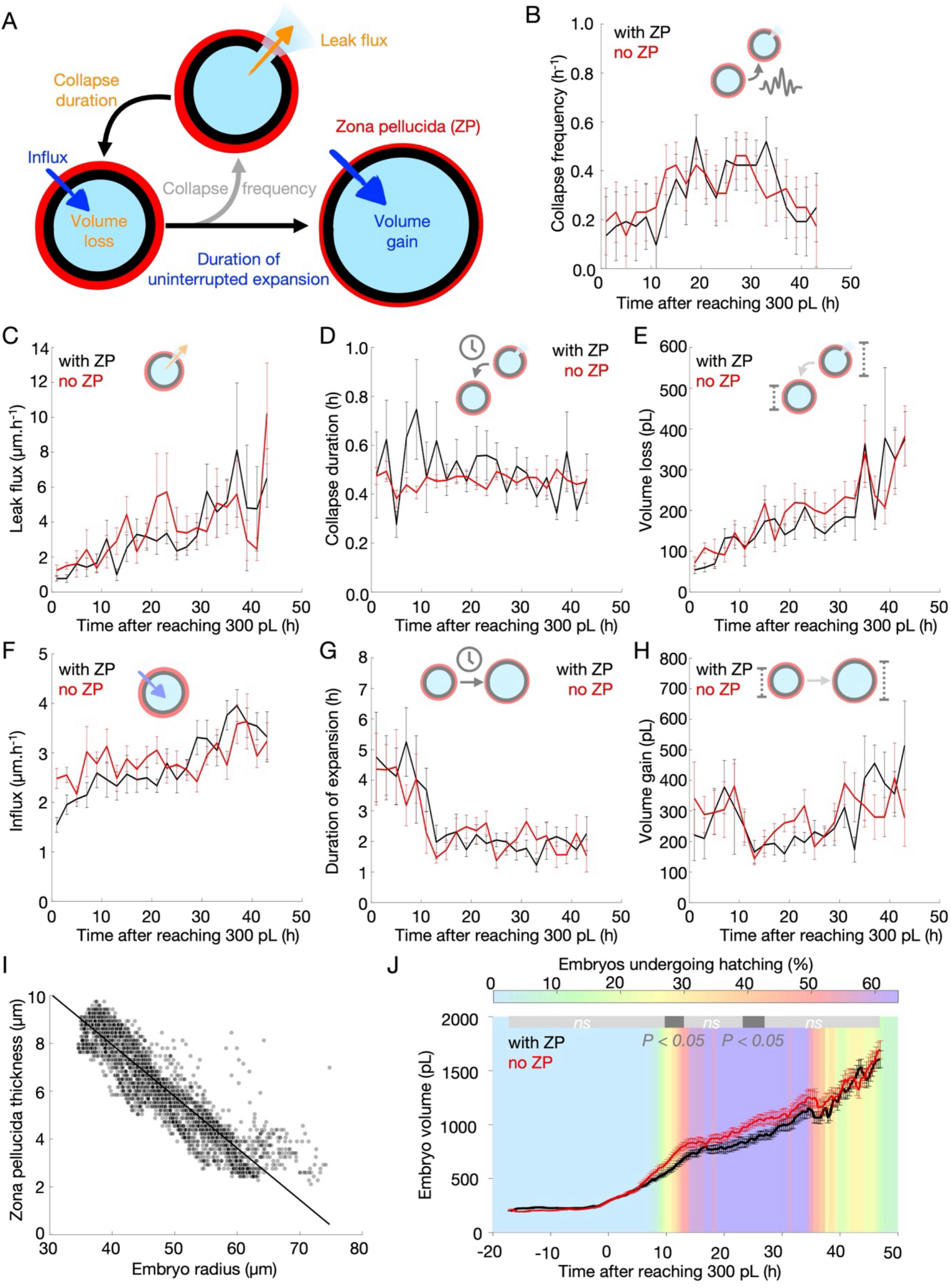
Analysis of distinct episodes during lumen expansion for embryos with and without zona pellucida reveals negligible influence of mechanical constraints on blastocyst expansion. **(A)** Schematics of the parameters extracted from individual embryos. **(B-H)** Collapse frequency (B), leak flux during collapse episode (C), duration of collapse episode (D), volume loss during collapse episode (E), influx during expansion episode (F), duration of expansion episode (G) and volume gain during expansion episode of 26 embryos from 3 replicates synchronized to the time they reached 300 pL. Mean ± SEM are shown for two-hour intervals. **(I)** Zona pellucida thickness as a function of embryo radius before they start hatching (Pearson correlation r = -0.96, with 3785 individual measurements on 26 embryos, Fisher’s test, p<10^−5^). **(J)** Volume of embryos with (black) and without ZP (red) aligned to the time they reached 300 pL (Mean ± SEM of 26 embryos with and without ZP each from 3 replicates, Mann-Whitney test result shown above the graph in light grey when not significant and in dark grey when p < 0.05). For embryos with ZP, the percentage of embryos that are hatching is color coded in the background.

Further investigating collapse dynamics over development, we measure that the rate at which fluid leaks out of the embryo accelerates between E3.5 and E4.5 (Fig 2C). With the duration of individual collapses staying stable at about 30 min long (Fig 2D), they are associated with larger and larger losses in volume over development, from 100 pL up to 400 pL per collapse between E3.5 and E4.5 (Fig 2E). Therefore, increased loss of volume across successive collapses can be explained by the accelerating leak flow rate (Fig 2C).

Increased overall leakage does not explain the continuous overall expansion of embryos (Fig 1C). To understand how embryos grow continuously despite increasing transient leakage flow rates, we analyzed the characteristics of episodes of uninterrupted expansion between individual collapses (Fig 2A). This reveals that fluid influx steadily increases over development with embryos growing from about 1.5 to 4 µm/h, corresponding to a flow rate of 1 to 4 pL/min (Fig 2F). Thanks to this, even with uninterrupted expansion phases shortening from 5 to 2 h (Fig 2G), the net gains between successive collapses stay between 200-400 pL throughout lumen expansion (Fig 2H). Therefore, overall lumen growth results from the steady increase in net fluid influx between collapses.

Taken together, our measurements show that embryos initially grow slowly but steadily due to modest fluid influx (Fig 1C and 2F). This uninterrupted inflation is followed by an overall slower expansion due to increased collapse frequency (Fig 2B) and leakage flow rates (Fig 2C) but expansion is maintained thanks to the ramping up of the fluid influx (Fig 2F).

### Mechanical constraint from the ZP marginally affects lumen expansion

Mechanical constraints were proposed to play a key role in regulating lumen growth in general and particularly in mouse blastocysts (Chan and Hiiragi, 2020). Specifically, mechanical stretch and elastic resistance of the epithelium were proposed to regulate the occurrence of collapses through mechanosensitive reinforcement of epithelial junctions (Chan et al., 2019). More generally, mechanical constraints from the embryo and/or its environment could influence the mechanical pressure of the lumen leading to increased leak flow rate during collapse (Lee et al., 2026) and slowed down fluid influx (Mosaliganti et al., 2019). While we observe increased leak flow rate during collapses (Fig 2C), we do not measure a slowdown in fluid influx but rather an increase (Fig 2F). To clarify the influence of mechanical constraints on expansion dynamics, we investigated the contribution of the ZP, an elastic material (Murayama et al., 2006; Yu Sun et al., 2003) controlling the mechanical pressure of embryos (Leonavicius et al., 2018).

As briefly mentioned, when embryos reach about 300 pL, they push against the ZP, which starts to thin (Fig 1B). Then, when the first collapse occurs, the ZP springs back to its original thickness (Fig 1B) and we calculate a very strong correlation between embryo volume and ZP thickness before hatching starts (Fig 2I). Consistent with this, when synchronizing and averaging embryos, we note a steady thinning of the ZP from about 9 to 5 µm until the beginning of hatching (Fig 1C). This indicates that the elastic shell closely follows the volume changes of the embryo, which, in turn, is mechanically compressed by the ZP, as measured previously (Leonavicius et al., 2018).

Given these multiple correlations between embryo expansion dynamics and ZP thickness, rupture and hatching, we directly tested the contribution of the ZP by removing it from embryos and analyzed their lumen formation dynamics (Fig 2J, Movie 2). Like control embryos, those without ZP showed two distinct expansion phases: first a fast inflation to 798 ± 34 pL within 12h after reaching 300 pL before adopting a slower expansion for the next 36 h (mean ± SEM, 26 embryos, Fig 1I). Therefore, despite showing very distinct mechanical properties (Chan et al., 2019; Leonavicius et al., 2018), embryos without ZP grow almost identically to control embryos. Looking into expansion dynamics in more details, we measure that removing the mechanical constraint provided by the ZP does not change leak frequency, rates, duration or net loss (Fig 2B-E). Embryos without ZP also show similar duration of uninterrupted expansion and net gain as control embryos (Fig 2G-H). The only measurable difference is a slightly faster influx during the initial uninterrupted expansion (Fig 2F), which does not have a long-lasting impact on embryo size since embryos with and without ZP reach the same size by 36 h after reaching 300 pL. Importantly, the remarkable change in expansion dynamics 12 h after reaching 300 pL, which coincides with the beginning of hatching for embryos with ZP, also takes place in embryos without ZP. In both embryos with and without ZP, this change in expansion dynamics is explained by the sudden increase in collapse frequency 12 h after reaching 300 pL (Fig 2B), which is also independent of the ZP. Therefore, changes in overall expansion dynamics and the frequency of collapses are independent from the mechanical constraints from the ZP and instead are due to properties that are intrinsic to the embryo.

Taken together, this analysis reveals that lumen expansion dynamics is only marginally affected by the presence of the ZP and its associated mechanical constraint. Therefore, lumen expansion dynamics are mainly set by the embryo itself rather than the mechanics of its environment.

### Dynamic osmotic shocks using microfluidics to study embryo transport properties

The prime determinants of blastocyst expansion dynamics are the fluid influx and the frequency of collapses (Fig 1-2), which are marginally affected by the mechanical constraints of the ZP (Fig 2). During uninterrupted expansion episodes, increased fluid influx could result from: 1/greater pumping of osmolytes, 2/lower permeability of the epithelium to osmolytes (e.g. increased tightness reducing their overall leakage after being actively transported) and/or 3/higher permeability to water allowing it to enter the embryo faster following changes in the osmotic gradient.

To investigate the transport properties of the blastocyst, we decided to override the osmotic transport of the embryo by acutely changing the osmolarity of the medium surrounding it. Since size changes after osmotic shocks usually occur within tens of seconds (Cadart et al., 2019; Roffay et al., 2021), we designed a microfluidic device allowing the hydrodynamic trapping of single embryos and the rapid change of medium with minimal perturbation while imaging embryos under a spinning disc confocal microscope (Fig 3A, Movie 3). To measure their size throughout osmotic shocks, we have developed an image analysis pipeline estimating embryo volume by segmenting its outline in the equatorial plane. Adding fluorescently labeled dextrans to media, we measure that the microfluidic chip allows medium exchange within less than 30 s while applying minimal hydrodynamic stresses onto embryos (Fig 2C, Fig S1, Movie 3-5).

**Figure 3:**
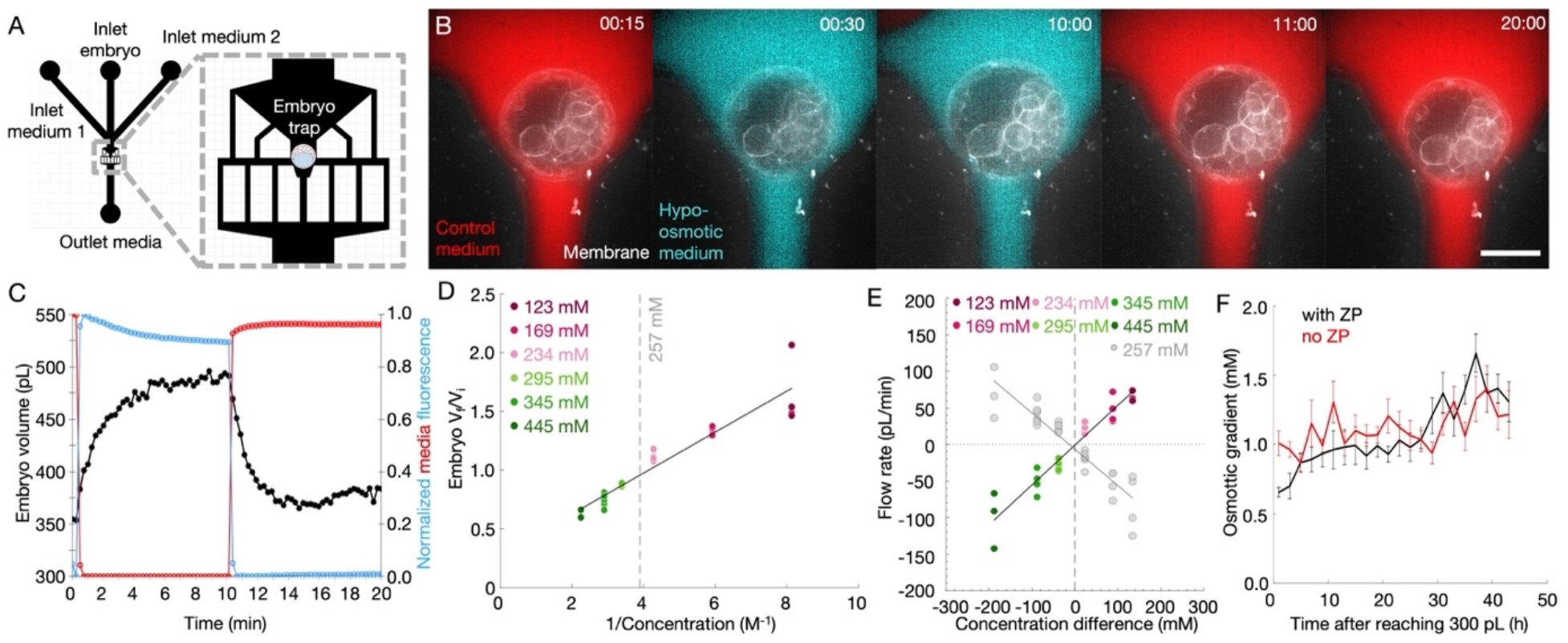
Osmotic pressure is sufficient to explain blastocyst size change and water transport rate. **(A)** Schematic of a microfluidic chip allowing dynamic control of embryo culture medium. Left: inlets are used to introduce different media and to load the embryo while the outlet at the bottom evacuates the media. Right: the main chamber is composed of microfabricated structures guiding the embryo to a central trap while maintaining parallel flows of media. **(B)** Maximum projection of confocal imaging of a blastocyst with fluorescent plasma membrane (grey) initially exposed to control medium (red, 257 mM) before the hypo-osmotic medium (blue, 169 mM) is pushed onto the embryo. The embryo swells before returning to its original size after reintroduction of the control medium. Time mm:ss, scale bar, 50 µm. **(C)** Embryo radius (black) during the changes between control (red) and hypo-osmotic (blue) media for the embryo shown in (B). **(D)** Boyle–van’t Hoff plot of blastocyst volume changes as function of the inverse concentration of hypo-osmotic (123 mM in purple, 169 mM in magenta and 234 mM in pink) and hyper-osmotic media (295 mM in pastel green, 345 mM in fern green and 445 mM in sea green). The grey dashed line indicates control medium concentration at 257 mM. Pearson correlation r = 0.96, 21 embryos, p < 10^−5^. **(E)** Volumetric flow rates for embryos subjected to osmotic shocks and recovery to control medium as a function of the difference in concentration between the exchanged media. Positive flow rates indicate accumulation of fluid into the embryo whereas negative values correspond to fluid leaving the embryo. Coloured dots correspond to the initial osmotic shocks (123 mM in purple, 169 mM in magenta, 234 mM in pink, 295 mM in pastel green, 345 mM in fern green and 445 mM in sea green) while grey dots show the recovery in control medium (257 mM). Osmotic shocks: linear fit gives y = - 0.57x + 143.94, Pearson correlation r = -0.98, 21 embryos, p < 10^−3^. Recovery: linear fit gives y = 0.46x - 119.55, Pearson correlation r = 0.95, 21 embryos, p < 10^−3^). **(F)** Osmotic gradient during physiological lumen growth of embryos with (black, 26 embryos from 3 replicates) and without ZP (red, 26 embryos from 3 replicates). The osmotic gradient is calculated from the linear scaling between flow rate and concentration presented in (F) and the physiological flow rate shown in Fig 2F.

Using microfluidics, we rapidly changed medium osmolarity from 257 mM to 123, 169 or 234 mM, using diluted medium, or 295, 345 or 445 mM, by adding sucrose. Importantly, this range of osmolarity is well beyond what embryos are suspected to encounter in vivo (Collins and Baltz, 1999). As expected, hyperosmotic media lead to the rapid deflation of embryos while hypoosmotic media cause them to rapidly inflate. In both cases, water flux following the shock slows down and reaches within 3-4 min an apparent plateau value (Fig 3C), suggestive of a new steady-state. Importantly, deflation of embryos after hyperosmotic shocks are different from collapses observed during development. Indeed, after hyperosmotic shocks only water leaves the embryo whereas during collapses fluid containing both water and osmolytes leak from the lumen. Collapses typically last half an hour (Fig 2D), presumably until the hole at the origin of the leak seals. In the case of osmotically-induced deflation, placing embryos back into normal medium allows them to gain back nearly all water that was lost within 3-4 min (Movie 4-5). Therefore, osmotic shocks appear reversible as long as no spontaneous collapse occurs. Together, these observations show that dynamic osmotic shocks using microfluidics permit measuring quantitatively the rates of passage of water through embryos to study their transport properties.

### Osmotic pressure rather than mechanical pressure sets blastocyst size

When subjected to osmotic changes, a semi-permeable membrane allows the passage of water, leading to a change in volumes of the two compartments it separates. When volume changes scale linearly with osmolarity, the system is called a perfect osmometer. This is represented as the volume ratio as a function of the inverse osmolarity following the so-called van’t Hoff plot (van’t Hoff, 1884; Olver and Benson, 2023). However, when the semi-permeable membrane is, for example, not infinitely stretchable, the volume change does not scale linearly with osmolarity since it would be limited by the mechanical pressure difference between the compartments (Cadart et al., 2019; Venkova et al., 2022). Therefore, to assess the transport properties of blastocysts, and in particular to test the potential influence of embryo mechanics, we used our microfluidic device to place embryos in media with different osmolarities.

Subjecting blastocysts to media with osmolarities ranging between 123 and 445 mM and measuring the ratio between their initial sizes and the one reached at steady-state gives the following van’t Hoff plot (Fig 3D). This reveals a linear scaling between embryo size change and osmolarity (Pearson correlation r = 0.96, 21 embryos, p < 10^-5^), indicating that over a broad range of osmolarity the blastocyst behaves as a perfect osmometer. This includes hypo-osmotic shocks leading to a rapid increase in blastocyst size against the mechanical resistance of the epithelium and of the ZP. Therefore, mechanical constraints of the embryos are negligible in front of osmotic changes within this range of concentration.

### Mechanical and transport properties co-evolve but do not significantly influence one another

To further test the influence of mechanical constraint during abrupt changes of volume due to osmotic shocks, we carried out similar osmotic shocks on embryos without ZP. Because loading embryos into the microfluidic chip tends to affect the integrity of embryos without ZP, we performed those shocks in culture dishes at the expense of dynamic analysis (Fig S2A). Imaging embryos before and 10 min after medium changes, we measured the volumes of embryos with or without ZP (Fig S2B). This revealed that, like control embryos, volume change of embryos without ZP scales linearly with osmolarity (Fig S2C). Therefore, the mechanical constraint of the ZP has no effect on the transport properties of embryos at steady-state.

Although most of the concentrations tested show a linear scaling, we note a slight deviation from linearity in the case of the lowest osmolarity in this dataset. This could be due to collapse after embryos rapidly gain a large amount of water, which are clearly noticeable when live-imaging on the microfluidic chip but more difficult without it. To assess this, we counted the number of embryos that would collapse when transferred into normal or hypotonic medium. Regardless of the presence of ZP or of the osmolarity, we note that ∼10% of embryos collapse upon medium transfer, even when transferred into media of the same tonicity (Fig S2D). Therefore, it is unlikely that deviation from linearity in this dataset would be due to collapses caused by sudden water influx. Instead, this further shows the importance of being able to follow dynamically the impact of osmotic changes.

Alternatively, osmolarity changes could affect embryo mechanical properties. To test the effect of rapid swelling after hypo-osmotic shocks on embryo mechanics, we measured blastocyst surface tension and mechanical pressure using microaspiration (Fig S3A). We measured that 10 min after being placed into a medium with reduced osmolarity of 169 mM, embryos increased in volume but their surface tension or mechanical pressure did not change consistently (Fig S3B-D). This is in contrast with the surface tension and mechanical pressure of embryos growing between E3.5 and E4.5, which increases (Fig S3E-F), in agreement with previous studies (Chan et al., 2019; Leonavicius et al., 2018). Therefore, acute swelling of the embryo and associated stretching of the epithelium is not necessarily associated with changes in tissue and lumen mechanics. Moreover, increased tension and mechanical pressure during development could result from changes other than the swelling of the lumen and stretching of the surface epithelium such as increased cell number or further differentiation of the TE.

To better understand the cellular changes associated with osmotic shocks, we used machine learning to segment the lumen from 3D stacks of transgenic embryos expressing a fluorescent membrane marker. This revealed that during osmotic shocks, both the luminal and cellular volumes of the embryo are affected (Fig S2E-F). Importantly, both cellular and luminal volume changes after osmotic shocks show a mostly linear scaling with medium concentration, indicating that both the cells and lumen behave like perfect osmometers (Fig S2C). Analyzing the scaling of volume adaptation to osmotic changes revealed that although both cells and lumens respond proportionally to the strength of osmotic shocks, the lumen contributes to most of the volume change both in absolute but also in relative values. Indeed, at all tested osmolarities, luminal volume change is ∼30% higher than its relative value compared to cellular volume change (Fig S2G). This indicates that, although osmotic shocks on blastocyst affect both cells and lumen, they primarily affect the lumen.

Taken together, these data show that osmotic pressure can dramatically change the size of blastocysts, without affecting embryo mechanics. Mechanical constraints do not seem to affect osmotically mediated blastocyst size adjustments whether directly by constraining embryo expansion or indirectly by inducing collapses. Therefore, the expansion of the embryo is dominated by osmotic rather than mechanical pressure.

### Water transport does not limit blastocyst dynamic size change

While blastocysts behave as perfect osmometers during osmotic shocks, their adaptation to the surrounding osmolarity is not instantaneous and depends on their transport properties. In particular, water permeability could limit the rate at which blastocysts change volume during osmotic shocks and potentially during physiological osmotic pumping in developing blastocysts.

To quantitatively assess this, we used our microfluidic chip to measure the rate at which blastocysts change volume during osmotic shocks (Fig 3B, Movie 4-5). During osmotic shocks, water flow rates can reach extremely high values up to 100 pL/min (Fig 3E). This is equivalent to half the initial volume of the embryo gained or lost within a minute and much higher than the influx and leakage rates measured during physiological expansion peaking at ∼3 and 20 pL/min respectively (based on Fig 2C and F). Despite such fast and large exchanges of water, the flow rates scale linearly with osmolarity (Fig 3E, Pearson correlation r = 0.98, 21 embryos, p < 10^-3^). This indicates that the water transport capacities of the embryos do not saturate within this range of osmolarities. Therefore, water transport during development is highly unlikely to be limiting the dynamics of blastocyst lumen expansion.

After osmotic shocks, normal medium is flushed back onto embryos, which return to their initial sizes (Fig 3B-C, Movie 4-5). Analyzing this recovery, we note flow rates that are identical to those measured during the initial shock (Fig 3E). This indicates that osmotic shocks and their associated mechanical stresses do not induce any detectable adaptation of the embryo. Therefore, despite extensive strains and strain rates, blastocysts do not appear to change their transport properties, or, as discussed before, their mechanical properties (Fig S3).

### Extraction of blastocyst transport properties from dynamic osmotic shocks

Given that transport properties dominate blastocyst expansion dynamics, we developed a theoretical model to quantify these transport properties from the dynamic response of embryos during osmotic shocks. The embryo is described as a spherical semi-permeable membrane containing water with a concentration *c*_*in*_ of a single type of electrically-neutral osmolyte (Fig 4A). The difference in concentrations between inside and outside the tissue creates a difference in osmotic pressure ΔΠ = k_B_T Δc = k_B_T(c_in_ – c_out_) with c_out_ the concentration of osmolytes in the medium outside, *T*, the temperature and *K*_*B*_ the Boltzmann constant. The tissue is assumed to be under tension, which induces a difference of Laplace pressure *ΔP = P*_*in*_ *– P*_*out*_ with *P*_*in,out*_ the mechanical pressures inside and outside the embryo, respectively. Using the framework of linear irreversible thermodynamics (Groot and Mazur, 2013; Onsager, 1931; Torres-Sánchez et al., 2021) and linearizing the chemical potentials, volume conservation and osmolyte conservation can be written as:

**Figure 4:**
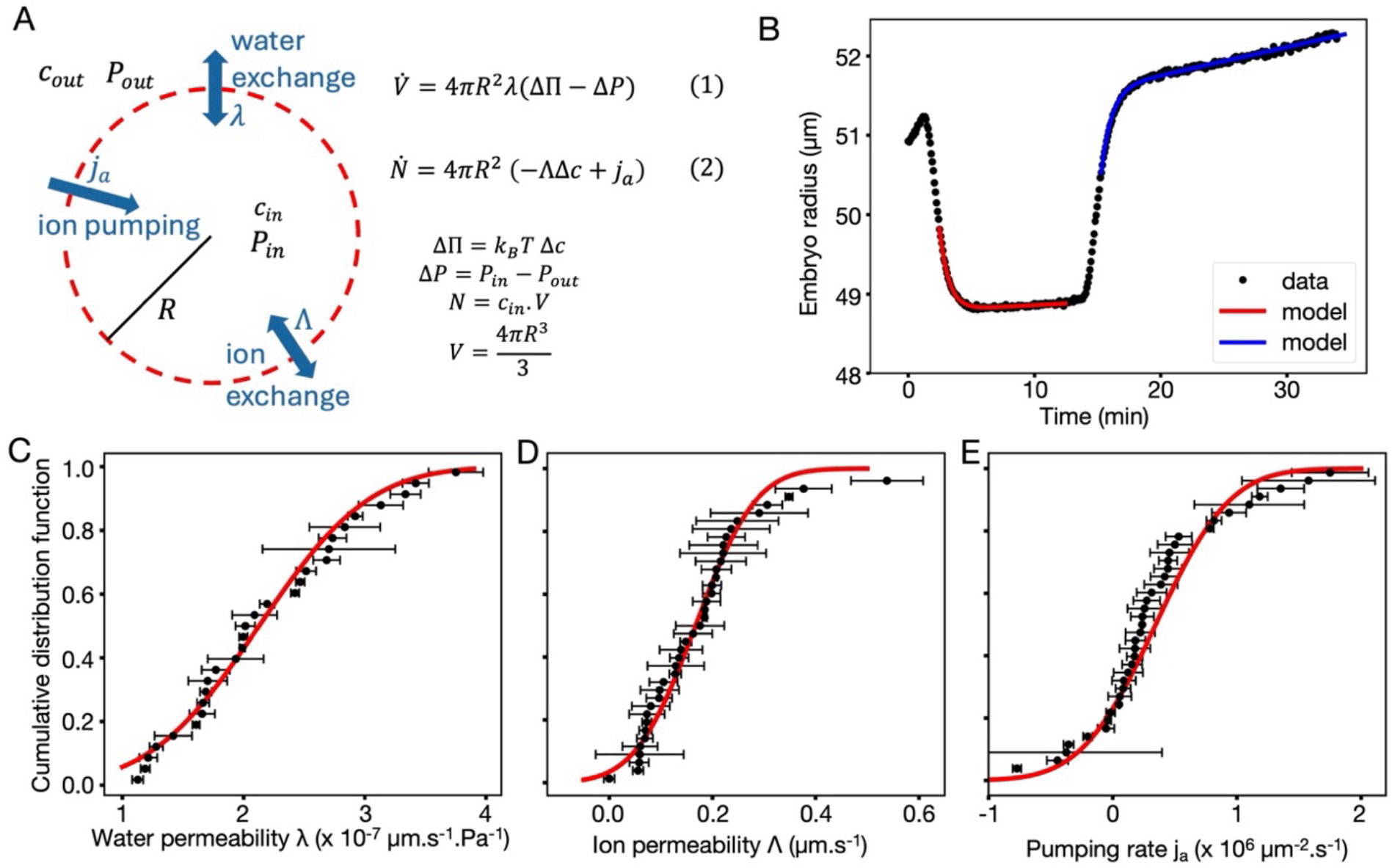
Theoretical model of embryo water transport. **(A)** Schematic diagram of a semipermeable membrane (red dashed line) changing radius R due to water exchange controlled by water and ion permeabilities, λ and Λ respectively, and ion pumping j_a_. c_out_, c_in_, P_out_ and P_in_ denote the external and internal concentrations and pressures respectively. Equations (1) and (2) use variations in embryo volume V and luminal osmolyte number N to extract water and ion permeability as well as ion pumping. Π is the osmotic pressure, k_B_ the Boltzmann constant, T the temperature, ΔP and Δc the pressure and concentration differences respectively. **(B)** Embryo radius over time during a hyper-osmotic shock experiment. Black dots correspond to the experimental data and the solid curves to model fits. **(C-E)** Cumulative distribution functions (CDFs) of the parameters obtained from the experiments: water (C) and ion permeabilities (D), and ion pumping (E). Each dot represents one shock; error bars indicate fitting uncertainty. Gaussian CDFs (red) are displayed for reference. *Guide:* in (d), ∼90% of λ values are ≤ 3 × 10^-7^ μm.s^-1^.Pa^-1^. Note that two outliers are not displayed on the graphs for readability (j_a_ = -2.0 ± 0.7 × 10^6^ μm^-2^.s^-1^ and Λ = 1.1 ± 0.2 μm.s^-1^; see corresponding data on Supp Theory Fig S4l and q, first curves.

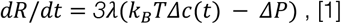

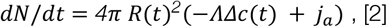

where *R(t)* is the radius of the embryo, *N(t)* is the number of osmolytes inside the lumen, γ and Λ are the permeabilities of the tissue to water and osmolytes, respectively, and *j*_*a*._ is the active pumping rate of osmolytes per unit area.

Considering embryos at steady-state after osmotic shocks, the mechanical pressure can be neglected compared to the osmotic pressure (Δ*P ≪* ΔΠ). Indeed, the mechanical pressure of the embryo is Δ*P* = 200-1000 Pa (Fig S3, Leonavicius et al., 2018; Chan et al., 2019; Dumortier et al., 2019), while for the osmotic shocks we applied, ΔΠ *≃ 10*^*5*^ *P*_a_= *≫* Δ*P* . In this case, Eq. [1] implies that the osmolyte concentrations inside the lumen and outside the epithelium must be equal at steady-state. This means *c*_*in*_ = *N*/(*V− V*_*d*_) =*c*_*out*_ with V = *4* π R^*3*^/*3* the volume of the embryo and *V*_*d*_ the volume of impermeant molecules. Considering that osmolyte exchange takes place on slower timescale compared to water transport, *N* is approximately constant during an osmotic shock. The steady-state equality of concentration can therefore be rewritten both for isotonic conditions (for which *V = V*^*iso*^ *and c*_*out*_ *= c*_*out*_^*iso*^) and for an arbitrary outer concentration, which yields the van’t Hoff relation:

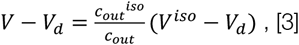

Beyond this steady-state description, Eqs. [1-2] allow us to accurately capture the dynamics of embryo size changes and to characterize its transport properties. For each embryo, the dynamics of the volume during osmotic shock experiments can thus be fitted (Fig 4B) to extract the pumping rate *j*_*a*_, the permeabilities to water λ, and to osmolyte Λ (Supp Theory). The distribution of those parameters is well-fitted by a normal distribution with mean and standard deviation of the experimental dataset (Fig 4C-E). This leads to a measurement of the water permeability *Λ = (2*.*1 ± 0*.*7) × 10*^*−7*^ *µm ⋅ Pa*^*−1*^ *⋅ s*^*−1*^, which matches previously reported values (Edashige et al., 2006; Leonavicius et al., 2018) and indicates that the embryo is rather permeable to water compared to other tissues (Timbs and Spring, 1996; Latorre et al., 2018; Torres-Sánchez et al., 2021). In addition, the embryo permeability to ions is found to be Λ = (0.17 ± 0.09) µm *⋅* s^−3^ and the active pumping rate is estimated as j_a_ = (3.4 ± 4.7) × 10^5^ µm^−6^ *⋅* s^−3^, which are both compatible with estimations of their orders of magnitude (Timbs and Spring, 1996; Latorre et al., 2018; Torres-Sánchez et al., 2021). Note that the model presented here considers permeabilities and pumping as constant per unit surface. We also considered the case in which the surface-integrated quantities of permeabilities and pumping are constant. The corresponding distribution of the fitted values are compatible with those reported here (Supp Theory). Therefore, our analysis cannot distinguish between permeabilities and pumping being constant per unit surface or being set per embryo. Both scenarios could explain the luminal fluid transport dynamics measured here.

The theoretical model and the parameters estimated from the osmotic shock experiments can then be used to revisit the inflation dynamics during embryo development (Fig 1C). First, we note that on this longer timescale, osmolyte dynamics can be considered quasistatic (*dN/dt* = *0*), such that the dynamics of the radius reduces to:

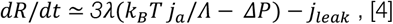

where we introduced a negative flux *j*_*leak*_ due to leaks. Let us first discuss the initial lumen growth phase during which collapses involving transiently increased leaks are rare (0-12 h after reaching 300 pL, Fig 1C). In that phase, *j*_*leak*_ is likely negligible, and the difference in expansion rate observed between embryos with and without ZP can be attributed to the mechanical constraint Δ*P*_*ZP*_ > *0* due to the ZP, which thus slightly but detectably slows down the expansion of embryos with ZP (Fig 2F). During the second phase (12 h after reaching 300 pL, Fig 1C), collapses become more frequent (Fig 2B) and leaks important *j*_*leak*_ > *0*, which reduces net embryo expansion rate. As embryos begin to hatch, the additional mechanical constraint due to the ZP disappears and the expansion rates of embryos with and without ZP becomes identical (Fig 2F).

Finally, assuming that osmotic pressure differences dominate over the mechanical pressure ones, Eq. [4] can be used to infer the osmotic pressure difference set by the embryo using the measured expansion rates between collapses (Fig 2F). This inference, which relies on the assumption that epithelial transport properties remain stable throughout development, yields an increase in osmotic pressure from ∼0.6 to 1.5 mM over 40 h (Fig 3F). Assuming removing the ZP does not change transport properties, we calculate a more modest increase from about 1 to 1.3 mM for embryos without ZP, which is consistent with their initially faster expansion compared to embryos with ZP.

Together, modeling and analysis of dynamic osmotic shocks allow extracting key transport properties of developing blastocysts, which increase the osmotic gradient with the outside medium to sustain their expansion despite more frequent collapses.

### Sodium transport by exchangers and pumps quantitatively determines osmotic pumping and lumen expansion

To build osmotic pressure between the outside medium and the lumen, blastocysts need to pump osmolytes and to have sufficiently low permeability to retain the pumped osmolytes. To quantitatively assess the role of ion pumps and exchangers in building osmotic pressure, we used ouabain and EIPA inhibitors of the sodium-potassium pump and sodium-proton exchangers respectively, which are known to regulate blastocyst lumen formation (Barr et al., 1998; Biggers et al., 1978; Dumortier et al., 2019; Kawagishi et al., 2004; Robinson et al., 1991; Schliffka et al., 2024; Wiley, 1984). As previously observed, treating E3.5 mouse embryos with ouabain or EIPA affects lumen expansion in a concentration dependent manner (Fig 5A). Synchronizing our analysis to the start of treatment, we find that 500 µM ouabain or 15 µM EIPA prevents inflation of embryos for over 15 h (Fig 5B). Analyzing the specific episodes leading to the lack of blastocyst expansion (Fig 5C-I), we find that fluid pumping rates are reduced from about to 2 to 1 or 0.5 pL/min between successive collapses when treated with 500 µM ouabain or 15 µM EIPA respectively (Fig 5G). Assuming that water permeability is not affected by the treatments, we infer that the osmotic pressure difference built by the embryo would be reduced from 1.5 mM to about 1 and 0.5 mM for 15 µM EIPA or 500 µM ouabain treated embryos respectively (Fig 5J). Also, we find that collapses are slightly less frequent with 500 µM ouabain or 15 µM EIPA (Fig 5C). However, since drug-treated embryos accumulate less fluid, leakages are smaller (Fig 5F) and detecting collapses becomes technically challenging, which could overestimate the effect of the drugs on this aspect.

**Figure 5:**
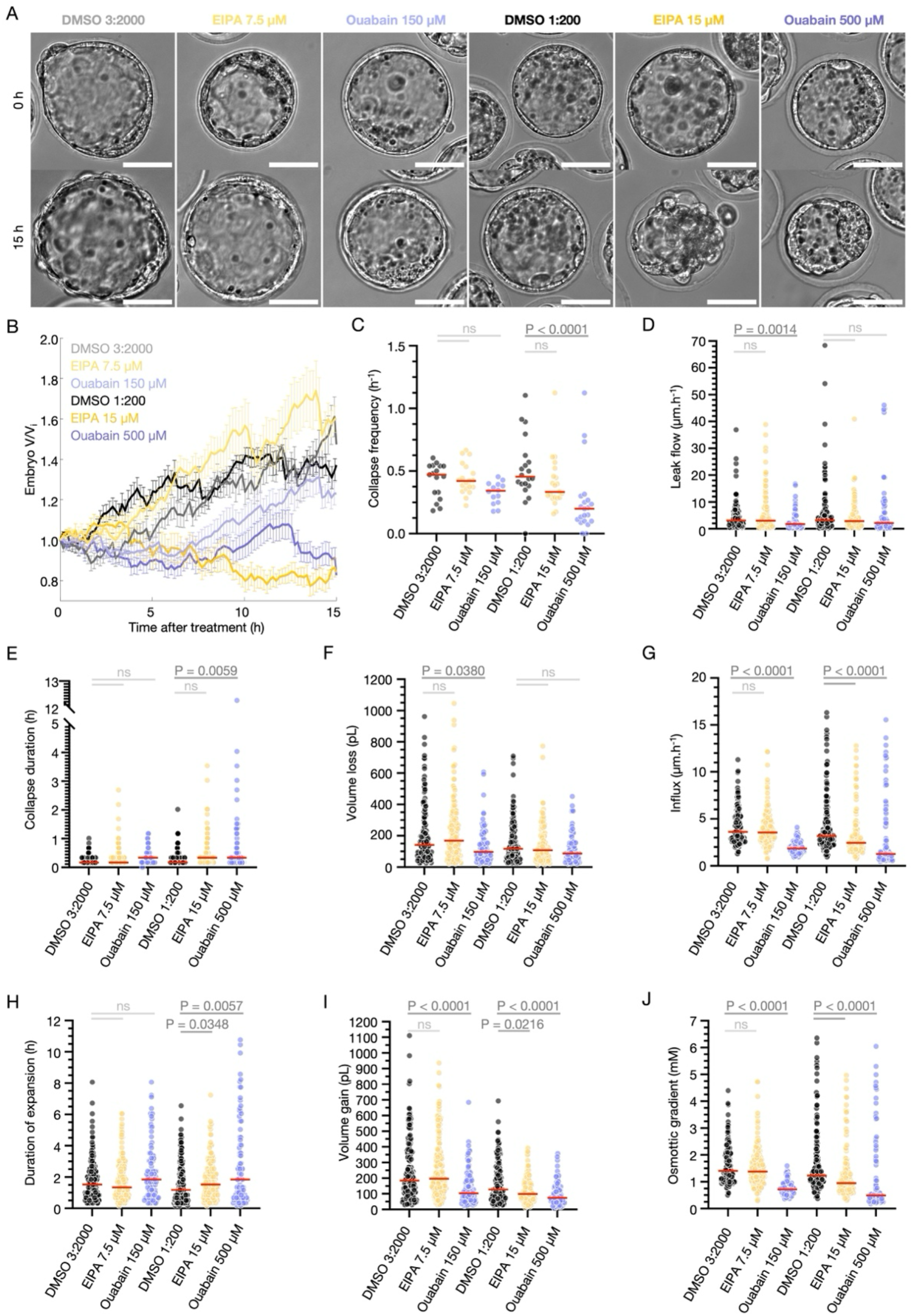
Ion transport sets the osmotic gradient controlling blastocyst expansion. **(A)** Brightfield images of E3.5 blastocysts before and after incubation with DMSO, EIPA or Ouabain. **(B)** Embryo volume V normalized to the initial volume V_i_ during treatment. Mean ± SEM of DMSO 3:2000 (grey, 17 embryos from 3 replicates), DMSO 1:200 (black, 20 embryos from 4 replicates), EIPA 7.5 µM (light yellow, 19 embryos from 3 replicates, Mann-Whitney tests non-significant over the entire period when compared with DMSO 3:2000), EIPA 15 µM (dark yellow, 21 embryos from 4 replicates, Mann-Whitney tests p < 0.05 from 5 h onwards when compared with DMSO 1:200), Ouabain 150 µM (light purple, 14 embryos from 3 replicates) and Ouabain 500 µM (dark purple, 21 embryos from 4 replicates, Mann-Whitney tests p < 0.05 from 3.5 h onwards when compared with DMSO). **(C-J)** Collapse frequency (C), leak flux during collapse episode (D), duration of collapse episode (E), volume loss during collapse episode (F), influx during expansion episode (G), duration of expansion episode (H), volume gain during expansion episode (I) and osmotic gradient (J) throughout 15 h after treatment with DMSO 3:2000 or 1:200, EIPA 7.5 or 15 µM or Ouabain 150 or 500 µM. P values from Kruskal-Wallis tests.

Together, our experiments and models allow determining the quantitative contributions of sodium-potassium pumps and sodium-proton exchangers to setting the osmotic pressure (Fig 5J). This confirms the essential role of osmotic pumping to draw water into the embryo during blastocyst formation. Since we found that water permeability does not limit fluid accumulation (Fig 3) and that mechanical constraints do not influence expansion dynamics (Fig 2), this makes osmotic pumping the main driver and regulator of blastocyst expansion.

## Discussion

The control of extracellular fluid transport is essential to many biological processes such as wound healing (Kennard et al., 2023), cell communication (Yoshiba et al., 2012; Faubel et al., 2016) or tissue shape and size in development (Barua et al., 2017; Mosaliganti et al., 2019; Yang et al., 2021; Stokkermans et al., 2022; Lee et al., 2026) and pathologies (Grubor et al., 2013; Shah et al., 2014). In this study, we investigated the regulation of fluid transport during the formation of the first mammalian lumen in the mouse blastocyst. Using time-lapse microscopy and image analysis, we find that this lumen initially grows fast when not interrupted by collapses and then continues growing at an overall slower pace when collapses become more frequent and massive. Importantly, we find that this change in dynamics is roughly concomitant with hatching but removing the ZP has little effect on blastocyst expansion two-phased dynamics. This indicates that hatching, and relief of associated mechanical constraints, does not trigger the increased collapse frequency slowing down overall blastocyst expansion. To sustain lumen inflation despite increased collapse frequency, we find that the rate of fluid accumulation between collapses increases over time. This larger fluid influx is explained by the ramping up of osmotic pumping rather than permeability to water since the speed of volume adaptation to artificial changes in osmolarity scales with a broad range of osmolarities. In fact, we develop a theoretical model to measure key transport properties such as water and ion permeability or ion pumping to understand what controls the inflation dynamics of the mouse blastocyst. In particular, we measure the osmotic pressure built by the embryo and find that it increases during development thanks to the pumping activity of cells. Together, we conclude that blastocyst expansion dynamics are primarily set by epithelial transport properties, which dominate mechanical constraints.

The implication of osmotic pumping and of epithelial transport properties in blastocyst expansion is not new. In fact, the idea that a material as complex as the blastocyst would behave as a perfect osmometer was proposed four decades ago when a few bovine embryos were subjected to different osmolarities at room temperature (Mazur and Schneider, 1986). Also, the roles of the active transporter Na/K ATPase or the passive Na/H exchangers, whose functions are inhibited by ouabain and EIPA respectively, in rabbit blastocyst formation were already identified half a century ago (Biggers et al., 1978; Powers et al., 1977). Finally, the osmolarity of the lumen of the rabbit blastocyst was measured on an osmometer after collection using a needle to puncture the embryo and it was found to be within a few mM to the one of the culture media event after varying media osmolarity (Borland et al., 1977). Yet, recent studies focused on the role of mechanical constraints of the environment (Leonavicius et al., 2018) or of the embryo itself (Chan et al., 2019) in controlling the size and expansion of the blastocyst without addressing the relative contributions of mechanics and transport properties. In this study, we have repeated some experiments carried out decades ago using modern quantitative approaches and we find that embryo expansion dynamics are dominated by osmotic transport rather than mechanical constraints. In the following paragraphs, we discuss how these and other studies can be brought into a consensual model of lumen expansion dynamics.

First, while recent studies proposed that hydraulic gating can lead to a steady-state lumen sizes (Ruiz-Herrero et al., 2017; Leonavicius et al., 2018; Chan et al., 2019; Mosaliganti et al., 2019), we report that embryos continuously grow and do not appear to reach a fixed size within at least 48 h after the start of lumen formation. However, the slowing down of overall expansion after the initiation of collapses could be mistaken for a steady-state if not imaged for sufficiently long and/or analysed with enough accuracy. Also, a steady-state size can be observed when embryos are cultured under strong artificial mechanical constraints (Leonavicius et al., 2018). Importantly, this steady-state size is small, as in (Chan et al., 2019), and does not seem to allow hatching, which is promoted by lumen expansion in vitro. Finally, with collapses spontaneously occurring every other hour followed by re-expansion, the notion of an embryo size that could be measured without considering expansion dynamics simply cannot be accurate. Therefore, previous experiments comparing embryos sizes without considering dynamic analysis of collapses should be interpreted with caution. Together, we conclude that, when cultured in conditions allowing normal development and hatching, there is no steady-state size for the mouse blastocyst.

Second, hydraulic gating implies that collapses are induced when increased tension in the epithelium would cause it to fail (Ruiz-Herrero et al., 2017; Chan et al., 2019; Mosaliganti et al., 2019). Collapses would thus become more frequent once embryos have reached a critical size, which would occur about 12 h after reaching 300 pL (Fig 1C). However, injection of fluid directly in the lumen (Chan et al., 2019) or hypo-osmotic shocks do not induce collapses (Fig S2E) despite increases in blastocyst surface area that are larger and faster than physiological ones (Fig 2F and 3E). This indicates that the blastocyst can cope with fast and large variations in surface stresses, which argues against hydraulic gating. A complementary cause of collapses is the occurrence of cell divisions in the epithelium (Leonavicius et al., 2018; Chan et al., 2019), which would cause transient increase in permeability due to the appearance of a new cell-cell contact with as of yet inefficient tight junctions. In fact, blocking cell division abolishes collapses (Chan et al., 2019). Cell divisions progressively become meta-synchronous during preimplantation development (Fujimori et al., 2009; Niwayama et al., 2019) and few of them can be observed in the hours directly following lumen opening. Occurring about 10 h later, the transition to the lumen expansion phase with higher frequency of collapses corresponds roughly to the duration of the cell cycle (Palmer and Kaldis, 2016; Abdelbaki et al., 2025). Moreover, the duration of collapses at about 30 min (Fig 2D) corresponds to the one of cytokinesis (Abdelbaki et al., 2025) and to the time needed for sister cells to build new junctions (Pinheiro et al., 2017). Why some divisions are associated with collapses while others are not remains to be determined but to our knowledge there is no evidence of collapse taking place in the absence of cell division. Together, cell divisions could temporarily increase permeability and increase the yielding threshold for the epithelium, thus triggering a cell division-dependent hydraulic gating.

Third, to allow for expansion beyond the yielding threshold, the hydraulic gating model would require this threshold to increase. Hence the addition to this model of contractility-dependent mechanosensing that would strengthen the epithelium and allow for further expansion. Indeed, as the blastocyst expands, mechanical pressure and surface tension increase (Leonavicius et al., 2018; Chan et al., 2019; Dumortier et al., 2019). Here, we confirmed this increase during physiological expansion but found little to no increase when artificially inflating blastocysts (Fig S3). Therefore, increased tension and mechanical pressure during development could result from changes other than the swelling of the lumen and stretching of the surface epithelium such as increased cell number or further differentiation and maturation of the TE. Furthermore, removing the ZP reduces mechanical constraints, associated mechanical pressure and surface stresses (Leonavicius et al., 2018), which should lead to reduced mechanosensitive reinforcement of the embryo (Chan et al., 2019). Yet, embryos without ZP show similar collapse frequencies to those with ZP. In our experiments consisting of osmotic shocks followed by recovery, we find that inflation and deflation dynamics do not change after abrupt manipulations of embryo size (Fig 3E). This suggests that embryos do not change their transport properties on the timescale of minutes in a detectable way despite rapid and large changes in lumen size. Finally, note that experiments aimed at perturbing contractility-dependent mechanosensing also affect cell divisions, which rely on contractility for cytokinesis (Schliffka et al., 2021). Therefore, disentangling contractility-dependent mechanosensing and cell division-dependent hydraulic gating can be difficult. Taken together, the contribution of mechanosensing to blastocyst expansion dynamics is unclear.

These observations lead us to propose the following model for blastocyst expansion dynamics. Shortly after finishing the 5^th^ wave of cleavage divisions and reaching the 32-cell stage, embryos start accumulating fluid. Until the 6th wave of cleavage divisions begins, lumen inflation is not interrupted by collapses, leading to fast expansion for about 10 h. When new divisions in the epithelium begin, transient increases in permeability cause leakage of luminal fluid and collapse for about 30 min until tight junctions are sealed back. Between collapses, inflation increases in speed and allows the embryo to reach larger and larger sizes. Note that this mechanism does not require mechanosensing to explain collapses nor does it need reinforcement of the epithelium to cope with higher stresses. This model is nonetheless compatible with the existence of mechanosensing.

Importantly, besides proposing an alternative explanation for lumen expansion dynamics that includes the relative contributions of epithelial transport properties and of tissue mechanics, our study provides key evidence for the dominant role of transport properties.

Inspired by early descriptions of the blastocyst behaving as a perfect osmometer (Mazur and Schneider, 1986), we verified this striking observation using live imaging of embryos in a microfluidic chip allowing dynamic media exchange. Indeed, despite its complexity, the blastocyst can be approximated to a spherical semi-permeable membrane whose volume is simply set by osmotic pressure with negligible influence from mechanical constraints. We verified this over a wide range of concentrations, 123-445 mM, which is likely exceeding the physiological variations in concentrations encountered by embryos in vivo, estimated between 200-300 mM (Menezo and Guerin, 1997; Collins and Baltz, 1999; Ismail et al., 2023), but remain relevant to embryo cryopreservation, which involves dehydration steps (Mazur and Schneider, 1986; Leonavicius et al., 2018; Mizobe et al., 2026). With such osmotic shocks, osmotic pressure vastly dominates the mechanical pressure of the embryo (ΔΠ ∼ 10^5^ >> ΔP = 200-1000 Pa) and the embryo behaves as a perfect osmometer. However, we estimate the osmotic gradient built by the embryo at ∼1 mM, which is in range with previous measurements performed at room temperature on rabbit embryos at ∼8 mM (Borland et al., 1977) or estimations from mouse embryos under strong mechanical constraints (Leonavicius et al., 2018). In the case of the osmotic gradient established by the embryo itself, the estimated osmotic and mechanical pressures are closer to one another (ΔΠ = 2000-4000 > ΔP = 200-1000 Pa), leaving the possibility for mechanical pressure to influence blastocyst inflation dynamics when osmotic pumping and/or permeability are weak. In fact, we estimate that the osmotic gradient increases during blastocyst maturation and we measure a small difference between the initial expansion dynamics of embryos with and without ZP. Overall, this effect is small and rapidly vanishes, indicating that transport properties dominate over mechanical ones.

Epithelial transport properties are much less well characterized than mechanical ones, especially in vivo (Lenne et al., 2021; Dagher et al., 2024; Lee et al., 2026). Combining modelling and microfluidics, we could estimate key epithelial transport properties in mouse blastocysts, namely ion permeability and active pumping, as well as water permeability. Our estimate of water permeability is in line with previous ones for the mouse embryo (Edashige et al., 2006; Leonavicius et al., 2018) and suggests a relatively high permeability to water compared to MDCK monolayers (Timbs and Spring, 1996; Latorre et al., 2018; Torres-Sánchez et al., 2021). This is consistent with the linear scaling of inflation and deflation rates during osmotic shocks. Even with ∼100 pL gained or lost per minute, the initially 200 pL mouse embryo can be traversed by water without slowing down. Therefore, water permeability is unlikely to be a limiting parameter when considering physiological dynamics of blastocyst expansion. Instead, inflation dynamics are regulated by the ability of the embryo to build a robust osmotic gradient. Our estimation of ion permeability, which reduces the ability to accumulate excess ions inside the lumen, is on par with the one made for MDCK monolayers (Fernández-Castelo et al., 1985). This leaves active ion pumping as the main driver to build the osmotic gradient that drives lumen expansion dynamics. We estimate active pumping rate at ∼10^5^ µm^-2^.s^-1^, which is in range with previous estimations in MDCK monolayers (Timbs and Spring, 1996; Latorre et al., 2018; Torres-Sánchez et al., 2021). Using drugs interfering with ion transport, we could quantify their effect on the osmotic gradient built by the embryo. Future studies, which could rely on the approaches developed here, will allow determining the relative contributions of ion transporters and pumps to the active pumping rates of the mouse embryo as well as other tissues.

Importantly, our model considers permeabilities to water and ions, and pumping rate, which are set constant per unit surface rather than constant for the whole embryo, an assumption that is commonly used in models of lumens (Dasgupta et al., 2018; Echeverría-Alar et al., 2026; Latorre et al., 2018; Shi et al., 2025; Torres-Sánchez et al., 2021; Wu et al., 2025). When surface area rapidly changes (e.g. collapses and osmotic shocks), one may question the validity of this assumption, since keeping these transport properties constant per unit surface would require fast channels and pumps turnover or activity changes. On the other hand, rapid changes in surface area of the epithelium could dilute or concentrate the proteins controlling epithelial transport properties and the maturation of the trophectoderm likely changes their compositions. Therefore, we also analyzed the osmotic shock data using constant surface-integrated quantities and obtained results which are also compatible with this scenario. In fact, our data cannot distinguish between these two hypotheses, and future experiments will be needed to disentangle these scenarios.

## Acknowledgements

We thank the Cell and Tissue Imaging Platform PICT-IBiSA of the Genetics and Developmental Biology Unit at Institut Curie (member of France-BioImaging ANR-24-INBS-0005 FBI BIOGEN) for their outstanding support; the animal facility of the Institut Curie for their invaluable help. We thank Anastasia Bekhtereva and Amaia Ochandorena-Saa for critical reading of the manuscript. L.D. is supported by the École Normale Supérieure via the Ecole Doctorale Frontières de l’Innovation en Recherche et Education - Programme Bettencourt and the Fondation ARC pour la recherche sur le cancer. Research in the labs headed by S.D. and J-.L.M. is supported by the Institut Curie, the Centre National de la Recherche Scientifique, Sorbonne Universite, Paris Sciences et Lettres and is funded by grants from the Agence Nationale de la Recherche (ANR-21-CE13-0027-01). The lab headed by S.D. is also funded by the ‘‘Institut Pierre-Gilles de Gennes’’ laboratoire d’excellence, ‘‘Investissements d’avenir’’ program ANR-10-IDEX-0001-02 and ITMO Cancer. The lab headed by J-.L.M. is also supported by the Institut National de la Sante et de la Recherche Médicale and by grants from the the European Research Council Consolidator Grant ERC-2022-CoG 101087656, the Coup d’elan from the Fondation Bettencourt Schueller and the European Molecular Biology Organization Young Investigator program.

## Author contributions

L.D. and L.d.P. performed experiments and prepared data for analyses with help from O.K.. L.D., G.G. and S.D. designed and optimized the microfluidic chip. L.D., C.C and A.M. designed and optimized the image segmentation pipelines. L.D., C.C., A.M., L.d.P and O.K. annotated images for segmentation model training. L.D., S.D. and J.-L.M. designed the project, analyzed the data. M.B. and C.D. developed the theoretical analysis. L.D., C.D., S.D. and J.-L.M. wrote the manuscript with inputs from all authors. L.D., C.D., S.D. and J.-L.M. acquired funding.

## Supplementary figures

**Supplementary figure 1:**
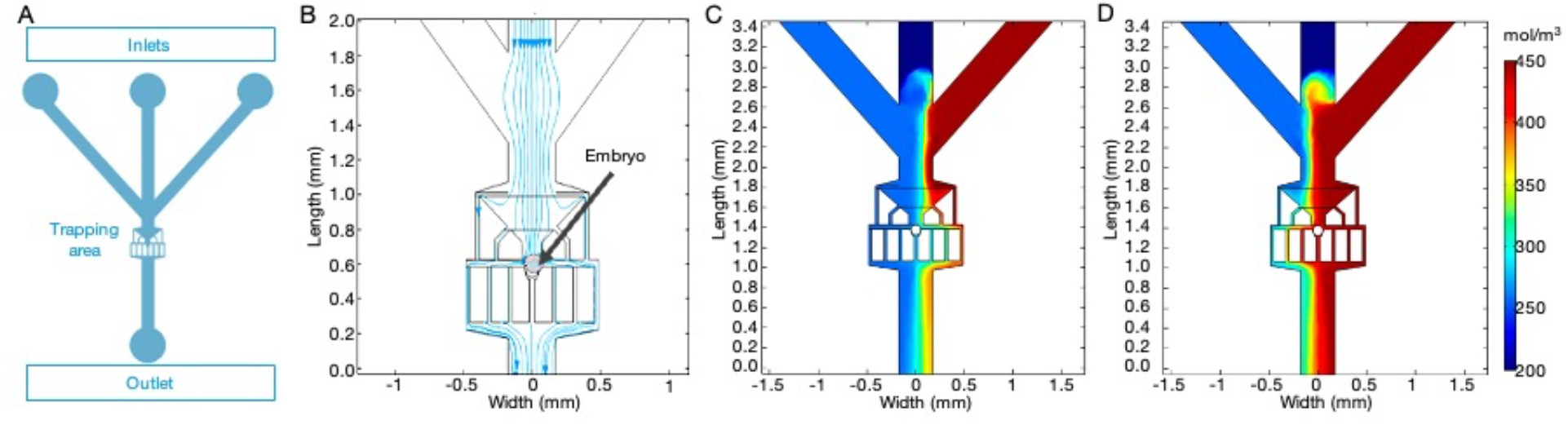
Microfluidic chip for embryo trapping and rapid medium exchange. **(A)** Schematic diagram of the microfluidic chip with inlet width at 350 µm, trapping width from 150 to 50 µm and lateral channels width at 30 µm. **(B)** COMSOL simulations of fluid streamlines showing the velocity field during embryo trapping with left and right inlets closed. **(C-D)** COMSOL simulations of medium exchange. Media from both left (257 mM) and right (455 mM) inlets flow into the chip and the position of the boundary between the different media is set by the respective pressures in the inlets. In (C), left inlet pressure is set to 16 mbar and the right inlet pressure to 5 mbar, setting the position of the boundary between media so that the embryo in the trap is exposed to the medium from the left inlet (257 mM). In (D), inverting the pressures of the left and right inlets moves the boundary so that the embryo in the trap is exposed to the medium from the right-hand side inlet (445 mM).

**Supplementary figure 2:**
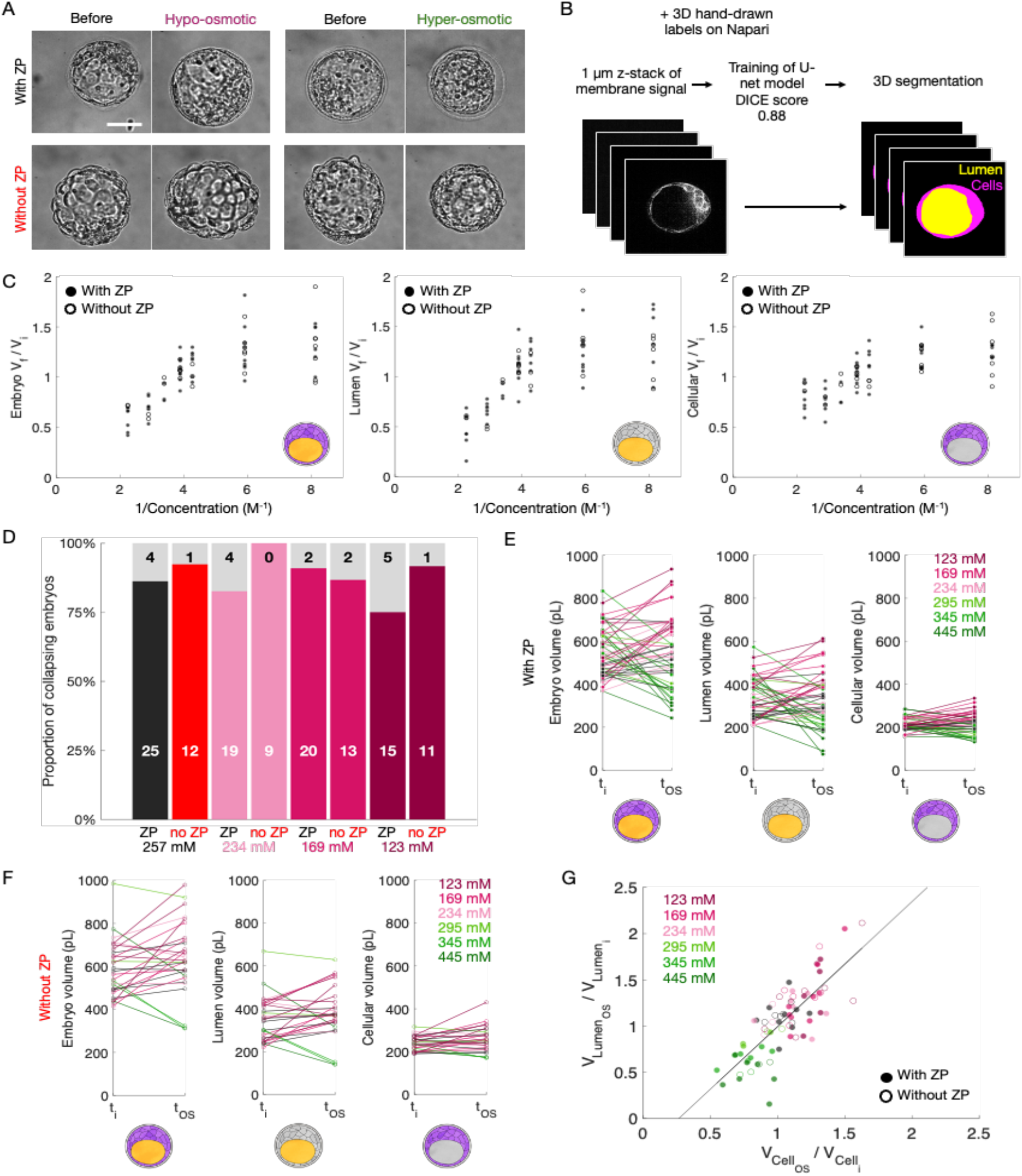
3D analysis of cellular and luminal contributions to volume changes at the steady-state of osmotic shocks on embryos with and without zona pellucida. **(A)** Images of embryos with (top) and without (bottom) zona pellucida (**ZP**) before and after 10 min in hypo-(left, 123 mM) or hyper-osmotic media (right, 445 mM). Scale bar, 50 µm. **(B)** Schematic of the segmentation of the cellular (magenta) and luminal (yellow) volumes of embryos with fluorescent membrane (left, grey). **(C)** Boyle–Van’t Hoff plots of embryos with (closed circles) and without **ZP** (open circles) giving their volume changes as a function of the inverse concentrations of hypo-osmotic (123, 169 and 234 mM) and hyper-osmotic media (295, 345 and 445 mM). The graphs show the volume change of the whole embryo (left), the lumen only (middle) or the cellular part of the embryo only (right) with respective Pearson correlations: r = 0.85, p < 10^-4^, r = 0.85, p < 10^-4^, r = 0.89, p < 10^-5^ for 49 embryos with **ZP** and 28 embryos without **ZP** from 8 replicates. **(D)** Proportion of embryos collapsing after hypo-osmotic shocks. The number of embryos is indicated within the bars (grey for collapsed embryos, black and coloured for non-collapsed embryos). **(E-F)** Individual volume measurements before and after osmotic shocks for embryos with (E, 49 embryos from 8 replicates) and without **ZP** (F, 28 embryos from 8 replicates). Whole embryo (left), luminal (middle) and cellular (right) volumes are provided. **(G)** Volume change for the lumen as a function to the one of the cells from embryos with (closed circles) and without **ZP** (open circles) subjected to osmotic shocks. Linear fit gives y = 1.33x - 0.36 indicating that luminal volume contributes proportionally 33% more to the total volume change compared to cells (Pearson correlation r = 0.86, p < 10^-5^,49 embryos with **ZP** and 28 embryos without **ZP** from 8 replicates).

**Supplementary figure 3:**
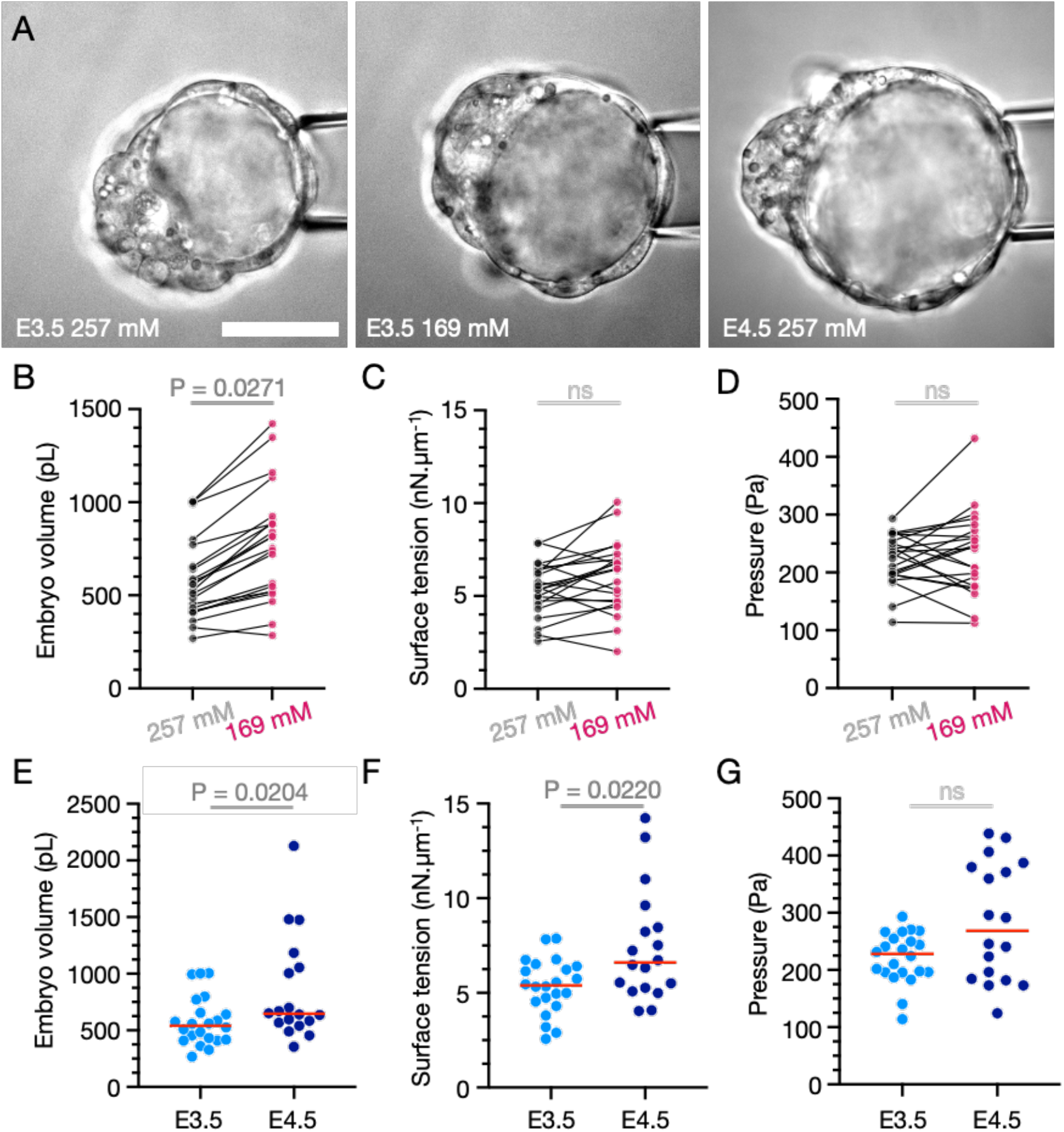
Mechanical characterization of blastocysts during lumen formation and after hypo-osmotic shocks. **(A)** Images of a blastocyst held by a micropipette at **E**3.5 in control (left) and hypo-osmotic (middle) media and at **E**4.5 in control medium (right). Scale bar, 50 µm. **(B-G)** Embryo volume (**B, E**), Surface tension (C, **F**) and mechanical pressure (**D**,G) for **E**3.5 embryos in control (257 mM) and hypo-osmotic (169 mM) media (**B-D**) and for blastocysts in control media at the **E**3.5 and **E**4.5 stages (**E-F**). Individual measurements for **B-D** before and after hypo-osmotic shocks for the same embryo are connected by a line.

## Supplementary movie legends

**Movie 1: Analysis of embryo and zona pellucida size during lumen formation and expansion**. Time lapse imaging of a mouse embryo with bright field images on the left and segmented embryo (cyan) and zona pellucida (ZP, red) on the right. Timepoint zero corresponds to the time at which the embryo reaches 300 pL for the first time. Before lumen formation, a series of cell divisions occur corresponding to the transition from 16- to 32-cell stage (-14 to -8 h). When fluid starts to accumulate within the embryo, cell-cell contacts show increased contrast and a small lumen appears (-7 to -5 h). The lumen increases in size bringing the embryo in contact with the ZP, which starts to thin (-5 to 0 h). Soon (5 h), first signs of ZP breaching appear and hatching completes about 30 h later. Throughout lumen expansion transient collapses can be observed. Images taken every 10 min, scale bar 50 µm.

**Movie 2: Analysis of embryo size during lumen formation and expansion without zona pellucida**. Time lapse imaging of a mouse embryo without zona pellucida (ZP) with bright field images on the left and segmented embryo (cyan) on the right. Timepoint zero corresponds to the time at which the embryo reaches 300 pL for the first time. Before lumen formation, a series of cell divisions occur corresponding to the transition from 16- to 32-cell stage (-20 to -4 h). When fluid starts to accumulate within the embryo, cell-cell contacts show increased contrast and a small lumen appears (-11 to -2 h). Throughout lumen expansion transient collapses can be observed. Images taken every 10 min, scale bar 50 µm.

**Movie 3: Dynamic medium change in a microfluidic chip**. Time lapse imaging of a blastocyst expressing a fluorescent membrane marker (grey) in red or cyan media. Since both media have the same osmolarity, the embryo size is not affected and only small movements of the embryo within the chip can be seen during media exchanges triggered by changes in hydrostatic pressure in the chip inlets. Brightfield shown on the left and fluorescent signal on the right. Images taken every 15 s, scale bar 20 µm.

**Movie 4: Hypo-osmotic shock and recovery in a microfluidic chip**. Time lapse imaging of a blastocyst expressing a fluorescent membrane marker (grey) in control medium (red) subjected to a 30% diluted medium (cyan). The embryo swells in the diluted medium and deflates when placed back in the control medium. Brightfield shown on the left and fluorescent signal on the right. Images taken every 15 s, scale bar 20 µm.

**Movie 5: Hyper-osmotic shock and recovery in a microfluidic chip**. Time lapse imaging of a blastocyst expressing a fluorescent membrane marker (grey) in control medium (red) subjected to a medium supplemented with 175 mM sucrose (cyan). The embryo deflates in the diluted medium without collapsing since it reinflates when placed back in the control medium. Brightfield shown on the left and fluorescent signal on the right. Images taken every 15 s, scale bar 20 µm.

**Movie 6-7: Analysis of embryo size during lumen formation and expansion in the presence of inhibitors of sodium transport**. Time lapse imaging of mouse embryos with bright field images annotated with embryos segmented in cyan. Timepoint zero corresponds to the time of addition of the treatment: DMSO, EIPA or Ouabain at mild (Movie 6) or high concentrations (Movie 7). Images taken every 10 min, scale bar 50 µm.

## Supplementary Theory

### I. MODEL FOR OSMOLYTE AND WATER TRANSPORT THROUGH AN EPITHELIUM

**FIG. S4:**
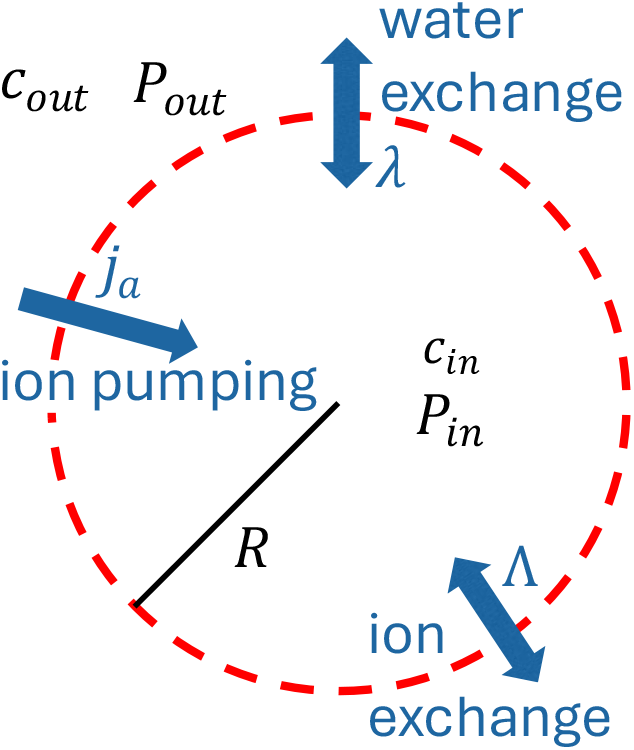
Schematic of the embryo as a semi-permeable membrane.

We developed a theoretical model to quantify the transport properties of the embryo and to analyze its dynamics during osmotic shock experiments. For completeness, We reproduce here the equations presented in the main text.

The embryo is described as a spherical semi-permeable membrane of radius *R* containing water with a concentration *c*_in_ of a single type of electrically-neutral osmolyte (see Fig S4). The di!erence in concentrations between inside and outside the tissue creates a di!erence in osmotic pressure Δ Π = *k*_*B*_*T* Δ*c* = *k*_*B*_*T* (*c*_in_ − *c*_out_) with *c*_out_ the concentration of osmolytes in the medium outside, *T* the temperature and *k*_*B*_ the Boltzmann constant. The tissue is assumed to be under tension, which induces a di!erence of Laplace pressure Δ*P* = *P*_in_ − *P*_out_ with *P*_in,out_ the mechanical pressures inside and outside the embryo, respectively. Using the framework of linear irreversible thermodynamics [1–3] and linearizing chemical potentials, volume conservation and osmolyte conservation can be written as:

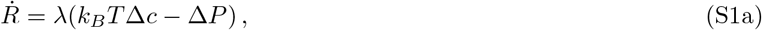

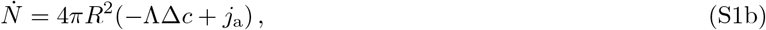

where 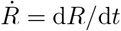 and where 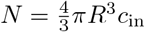is the number of osmolytes inside the membrane, *λ* the water permeability and Λ the ion permeability. It is also assumed that the embryo actively pumps osmolytes at a constant rate *j*_a_ per unit area.

### B. Dimensionless equations

In the following, we consider that *j*_a_ and Δ*P* vary suffciently slowly to be considered constant over a single shock. Under this assumption, Eq. (S1) reduce to a single equation for the dimensionless radius *x* = *R/R*_0_ with *R*_0_ = *R*(*t* = 0) that reads:

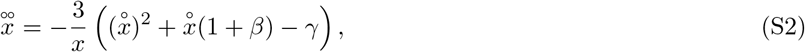

where we introduced the dimensionless time derivative 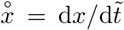 with 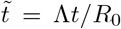 the dimensionless time. The dimensionless parameters ***β*** and *γ* are defined as:

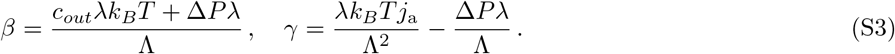

**FIG. S5:**
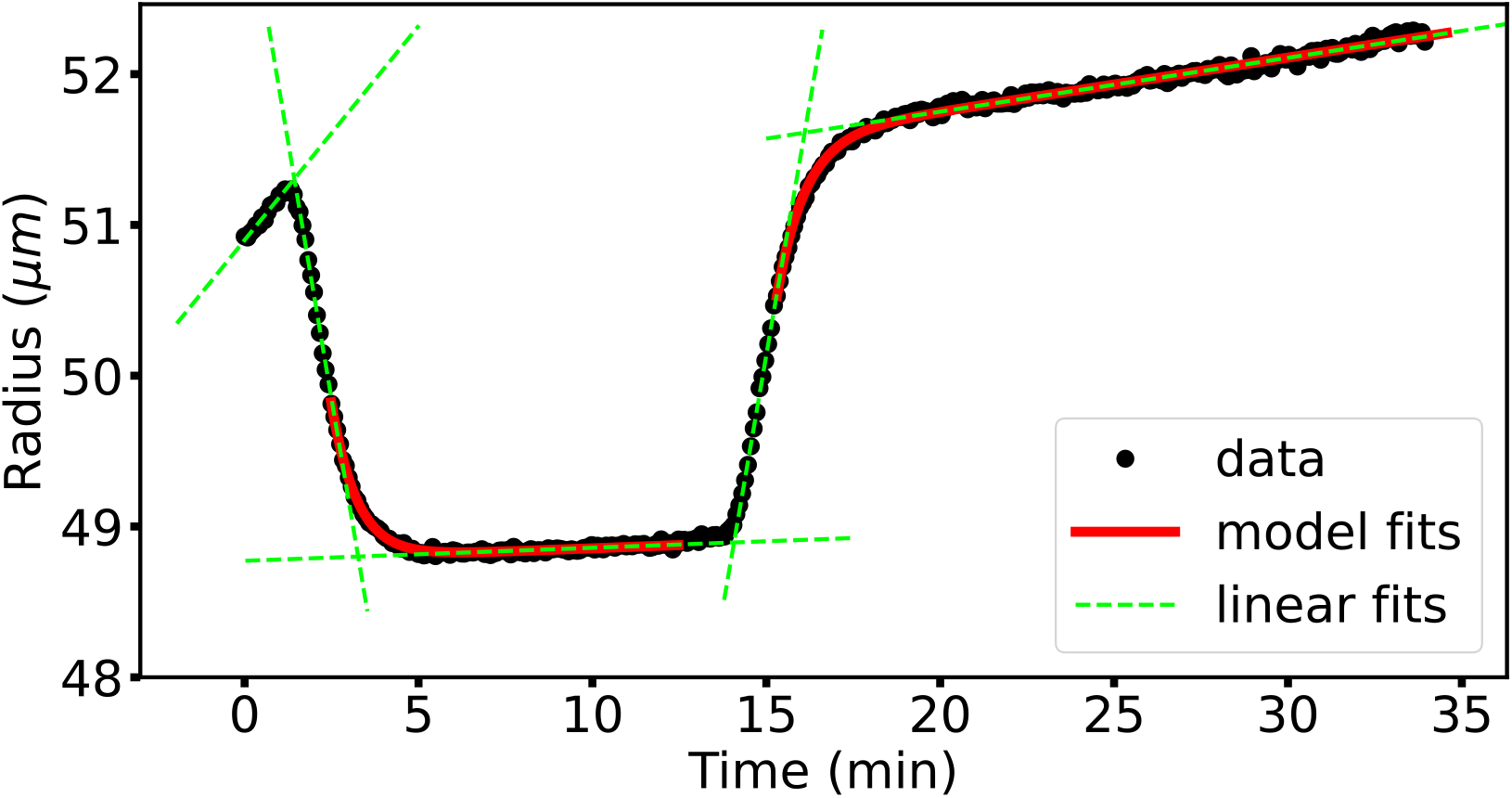
Osmotic shock response of a blastocyst. Osmotic shocks were performed on a blastocyst, starting with a solution of ions at a total concentration of 257 mM. The concentration was increased up to 300 mM at *t* = 1.2 min (forward shock), and then brought back to 257 mM at *t* = 14.1 min (backward shock). Black dots: experimental data. Dashed green lines: linear fits of the dynamics before and after each shock. Solid red lines: fit of the model described in Eqs. (S1) and (S2).

We note that since a concentration cannot become negative, Eq. (S2) directly implies that the speed 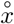 has a minimum: 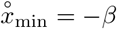. Thus, only one of the two roots of the polynomial in Eq. (S2) is a physically-acceptable stationary solution, which reads:

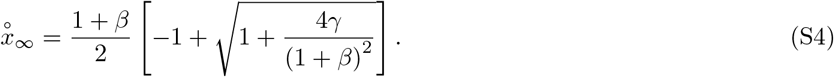

## II. FIT OF THE DYNAMIC RESPONSE TO OSMOTIC SHOCKS

### A. Water permeability of the embryo

In order to fit the model to the data, two subsequent steps are followed. First, we fitted the rate of increase of the radius right before and after the shock (green lines in Fig S5). Indeed, this allows for a measure of the water permeability *ω* using the relation:

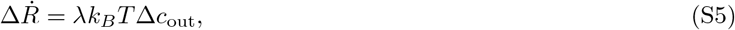

where 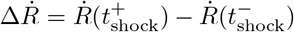 is the difference in the radius growth rate after and before the shock, and 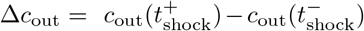 is the difference in concentrations outside after and before the shock. Relation (S5) is plotted on Fig S6a using the experimental data for both the forward and backward osmotic shocks. Rather than using Fig S6a to determine an average permeability, we extracted one value per shock, as different embryos could have different permeabilities. Figure 4C in the main text displays the corresponding cumulative distribution function of the permeability 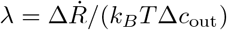. In 10 out of 21 experiments, the recording only starts at the instant of the first shock. No individual permeability can be inferred for forward shocks in these experiments, so they are not included in Figure 4C. However, in order to infer Λ and *j*_a_, these shocks are assigned the average global permeability with a standard error equal to the standard deviation of the normal distribution displayed on Figure 4C.

**FIG. S6:**
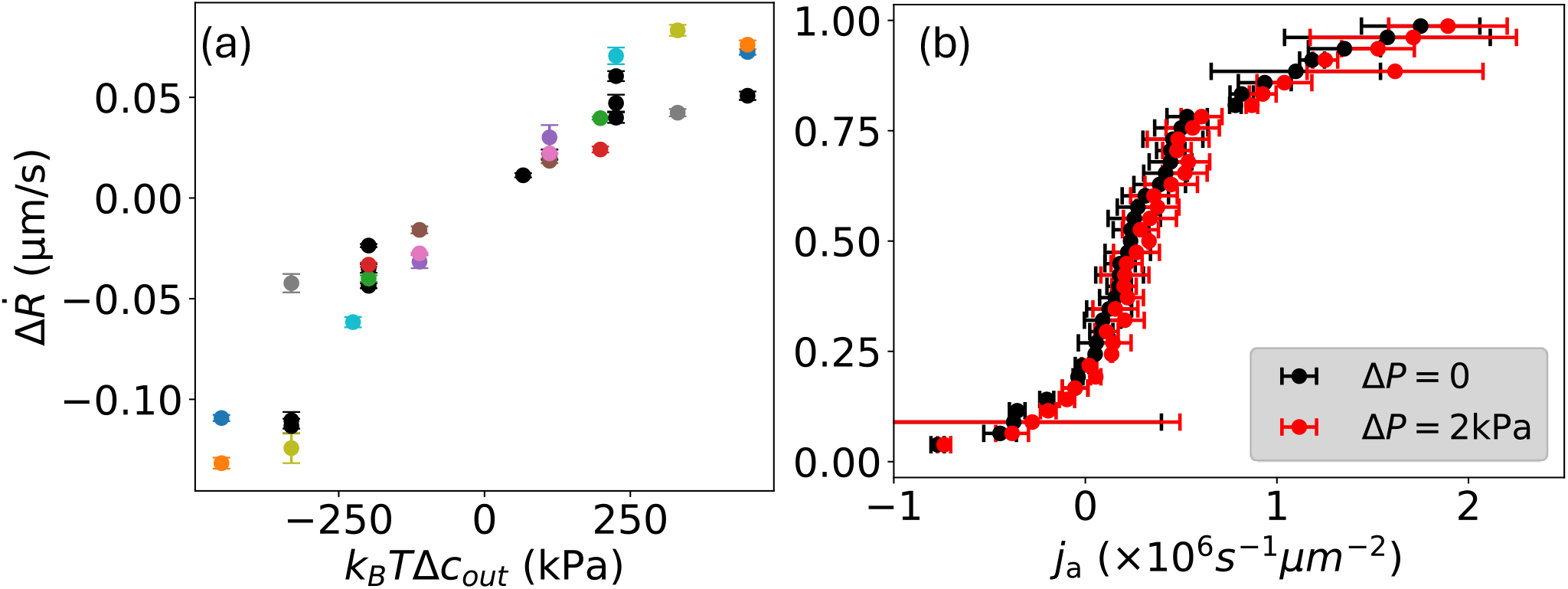
Parameters extracted from the model. (a) Difference in velocity after and before the shock 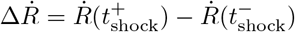 as a function of the concentration difference 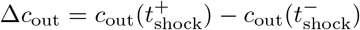. Black dots correspond to experiments where the slope before the forward shock was not recorded and therefore no permeability was computed. Coloured dots correspond to experiments where backward and forward permeabilities were computed (2 points). Corresponding panels in Fig S7: 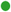 (h) 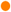 (u) 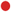 (i) 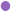 (k) 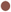 (m) 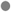 (c) 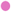(n) 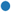(s) 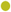(d) 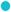(b). Cumulative distribution function of the pumping rate *j*_*a*_ obtained from fitting the experimental data and considering a pressure difference Δ*P* = 0 (black dots). Corresponding curve when a pressure difference Δ*P* = 2 kPa (red dots) is considered.

### B. Osmolyte permeability of the embryo

In order to obtain the osmolyte permeability, we first rewrite Eq. (S2) as:

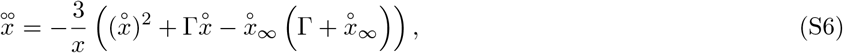

where Γ = (1 + *β*) and where we introduced the (dimensionless) stationary rate of increase of the radius 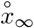 defined by 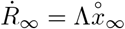. The dimensionless form (S6) is better suited for fitting since the value of 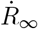 can be measured from experimental data as the stationary slope after the shock. Therefore, a single dimensioless parameter, Γ, remains to be fitted in Eq. (S6).

Note that we considered in our computations that *c*_out_ varied as a step function of time while in the experiments the concentration change takes a time of the order of a minute. To account for this simplification, we introduce an additional parameter *t*_0_, the effective shock time. Therefore, we eventually fit Eq. (S6) with two parameters: Γ and *t*_0_, and typical fits (one for the forward and one for the backward shock) are displayed on Fig S5. We also provide the result of the fitting procedure for all experiments in Fig S7.

In detail, once the initial (respectively final) rate of increase 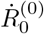 and its error *σ*_0_ 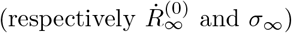 are determined by fitting the asymptotes, the fitting of the nonlinear model is done by minimizing the following function, jointly on & and *t*_0_, and the initial and final rates of increase 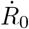 and 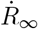:

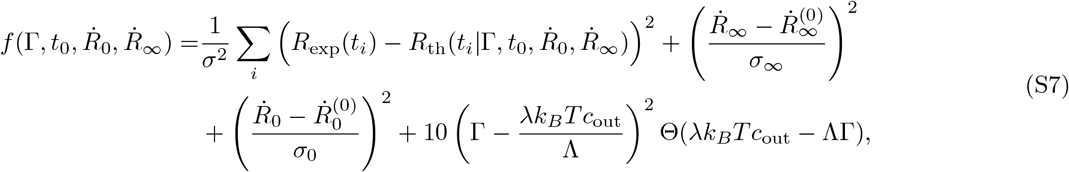

with Θ (*z <* 0) = 0 and Θ (*z ≥* 0) = 1, and where *σ* = 0.1 *µ*m is the typical standard error. The last term in Eq. (S7) enforces that Λ remains non-negative.

**TABLE S1:** Summary of the results displayed on Fig 3 of the main text.

| Quantity | Value |
| --- | --- |
| $\lambda$ ($10^{-7} \mu\text{m} \cdot \text{s}^{-1} \cdot \text{Pa}^{-1}$) | $2.1 \pm 0.7$ |
| $\Lambda$ ($\mu\text{m} \cdot \text{s}^{-1}$) | $0.17 \pm 0.09$ |
| $j_a$ ($10^5 \text{s}^{-1} \cdot \mu\text{m}^{-2}$) | $3.4 \pm 4.7$ |

One notices at this point that three physical parameters (Λ, Δ*P* and *j*_a_) must be extracted, while only two dimensionless parameters 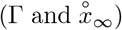 are obtained from our fitting procedure, which indicates that determining them separately may *a priori* not be possible. Some progress can nonetheless be made by analysing typical orders of magnitude. Indeed, the mechanical pressure difference Δ*P* is of the order of 500 Pa (Supplementary Figure 3 and Refs. [4–6]), while for the osmotic shocks analysed here, *k*_*B*_*Tc*_out_ ≃6 x 10^5^ Pa ≫ Δ*P* . Neglecting the contribution of Δ*P* in Γ thus yields:

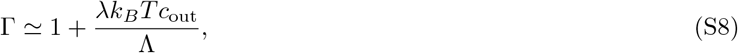

and allows for a measurement of the osmolyte permeability Λ using the already measured value of the water permeability *λ*. The corresponding cumulative distribution function of the values measured on the curves is displayed on Fig 3D of the main text.

### C. Active osmolyte pumping rate

The evaluation of the pumping rate *j*_a_ is more difficult as it cannot be separated from the pressure term in 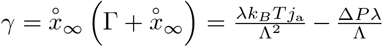. To make progress, we considered two extreme values for the mechanical pressure difference (Δ*P* = 0 and Δ*P* = 2 kPa), and computed the corresponding value of the pumping rate *j*_a_ for those two values, see Fig S6b. We observe that these two extreme values provide two pumping rate distributions which are almost identical, and thus allow us for a relatively accurate estimation of the pumping rate. We therefore obtain a value for *j*_a_ that ranges between 10^5^ and 10^6^ s^*−*1^ · *µ*m^*−*2^, which is surprisingly close to the order-of-magnitude estimate provided in Ref. [3].

Interestingly, some values of the pumping rate *j*_a_ seem to be negative. These negative values are mainly due to the fact that a few embryos display a negative steady-state growth rate following a hyperosmotic shock (see Fig S7). This may suggest that some shocks, especially the strongest, might affect the effective pumping of the embryo. However, it is worth noticing that the “normal” (inward pumping) phenotype is recovered after a backward shock.

Table S1 recapitulates all the dimensionful quantities estimated from the osmotic shock experiment in the blastocyst.

---

**FIG. S7:**
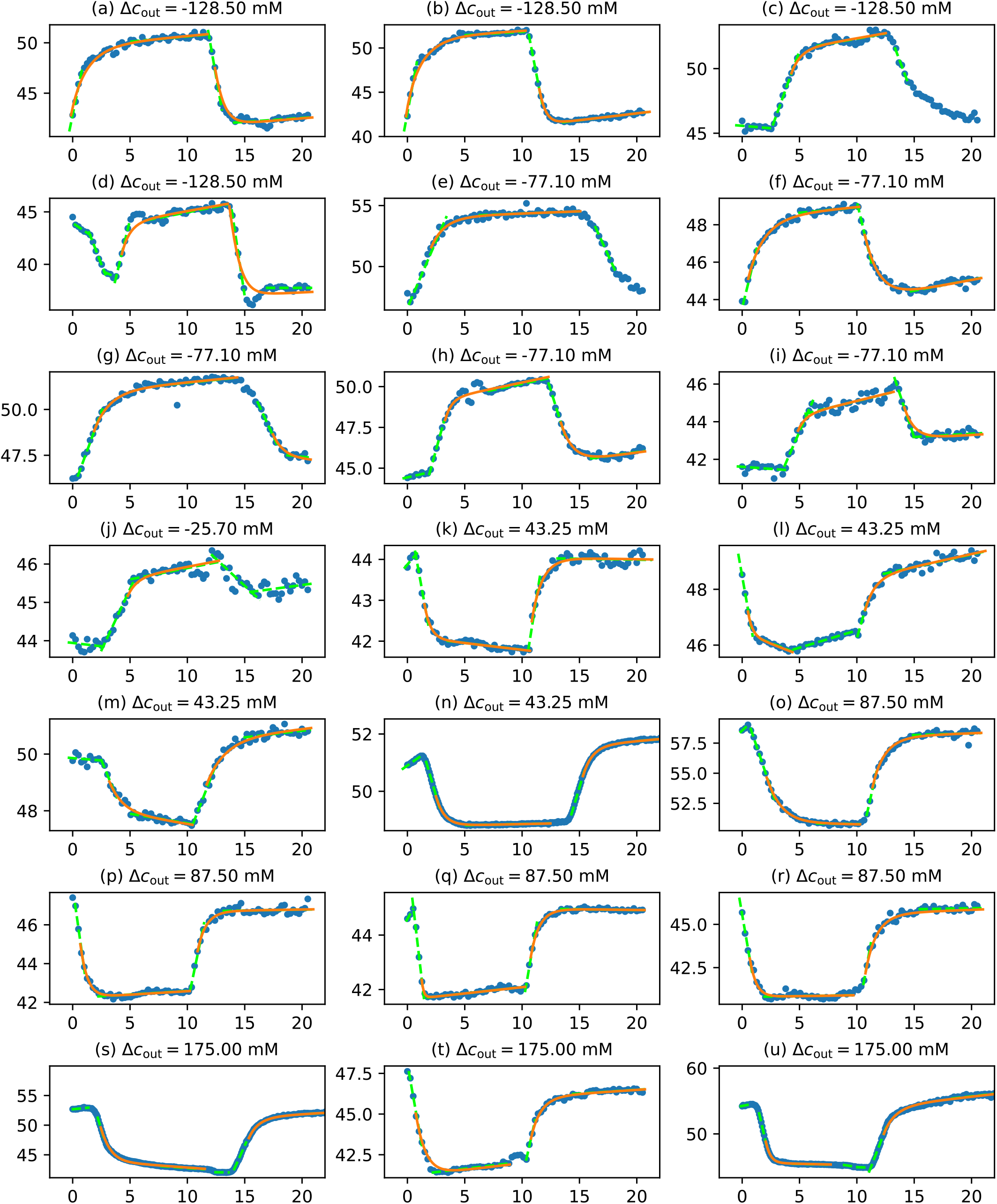
Experimental curves and corresponding fits. Radius of the embryo (*µ*m) as a function of time (min) from the experiments (blue dots) and corresponding fit by the model (solid orange lines). Green dotted lines correspond to linear fits. Titles give the extra concentration that was added (or removed) on the first shock.

## References

Abdelbaki, A., McCarthy, A., Karsa, A., Muresan, L., Elder, K., Papathanasiou, A., Snell, P., Christie, L., Wilding, M., Steventon, B. J., et al. (2025). Live imaging of late-stage preimplantation human embryos reveals de novo mitotic errors. Nat Biotechnol.

Barr, K. J., Garrill, A., Jones, D. H., Orlowski, J. and Kidder, G. M. (1998). Contributions of Na+/H+ exchanger isoforms to preimplantation development of the mouse. Mol. Reprod. Dev. 50, 146–153.

Barua, D., Parent, S. E. and Winklbauer, R. (2017). Mechanics of Fluid-Filled Interstitial Gaps. II. Gap Characteristics in Xenopus Embryonic Ectoderm. Biophysical journal 113, 923–936.

Biggers, J. D., Borland, R. M. and Lechene, C. P. (1978). Ouabain-sensitive fluid accumulation and ion transport by rabbit blastocysts. The Journal of physiology 280, 319–330.

Biggers, J. D., McGinnis, L. K. and Raffin, M. (2000). Amino Acids and Preimplantation Development of the Mouse in Protein-Free Potassium Simplex Optimized Medium. Biology of reproduction.

Borland, R. M., Biggers, J. D. and Lechene, C. P. (1977). Fluid transport by rabbit preimplantation blastocysts in vitro. Reproduction 51, 131–135.

Cadart, C., Venkova, L., Recho, P., Lagomarsino, M. C. and Piel, M. (2019). The physics of cell-size regulation across timescales. Nat. Phys. 15, 993–1004.

Chan, C. J. and Hiiragi, T. (2020). Integration of luminal pressure and signalling in tissue self-organization. Development 147, dev181297.

Chan, C. J., Costanzo, M., Ruiz-Herrero, T., Mönke, G., Petrie, R. J., Bergert, M., Diz-Muñoz, A., Mahadevan, L. and Hiiragi, T. (2019). Hydraulic control of mammalian embryo size and cell fate. Nature 571, 112–116.

Collins, J. L. and Baltz, J. M. (1999). Estimates of Mouse Oviductal Fluid Tonicity Based on Osmotic Responses of Embryos1. Biology of Reproduction 60, 1188–1193.

Dagher, L., Descroix, S. and Maître, J.-L. (2024). Intercellular fluid dynamics in tissue morphogenesis. Current Biology 34, R1031–R1044.

Dasgupta, S., Gupta, K., Zhang, Y., Viasnoff, V. and Prost, J. (2018). Physics of lumen growth. Proceedings of the National Academy of Sciences of the United States of America 115, E4751–E4757.

Dumortier, J. G., Le Verge-Serandour, M., Tortorelli, A. F., Mielke, A., de Plater, L., Turlier, H. and Maître, J.-L. (2019). Hydraulic fracturing and active coarsening position the lumen of the mouse blastocyst. Science 365, 465–468.

Echeverría-Alar, S., Narasimhan, B. N., Fraley, S. I. and Rappel, W.-J. (2026). Lumens as active balloons: a biological physics review.

Edashige, K., Tanaka, M., Ichimaru, N., Ota, S., Yazawa, K., Higashino, Y., Sakamoto, M., Yamaji, Y., Kuwano, T., Valdez, D. M., Jr., et al. (2006). Channel-Dependent Permeation of Water and Glycerol in Mouse Morulae1. Biology of reproduction 74, 625–632.

Faubel, R., Westendorf, C., Bodenschatz, E. and Eichele, G. (2016). Cilia-based flow network in the brain ventricles. Science (New York, NY) 353, 176–178.

Fernández-Castelo, S., Bolívar, J. J., López-Vancell, R., Beaty, G. and Cereijido, M. (1985). Ion Transport in MDCK Cells. In Tissue Culture of Epithelial Cells (ed. Taub, M.), pp. 37–50. Boston, MA: Springer US.

Firmin, J. and Maître, J.-L. (2021). Morphogenesis of the human preimplantation embryo: bringing mechanics to the clinics. Seminars in Cell & Developmental Biology S1084952121001920.

Fujimori, T., Kurotaki, Y., Komatsu, K. and Nabeshima, Y. (2009). Morphological organization of the mouse preimplantation embryo. Reproductive sciences (Thousand Oaks, Calif.) 16, 171–177.

Fütterer, C., Colombo, C., Jülicher, F. and Ott, A. (2003). Morphogenetic oscillations during symmetry breaking of regenerating Hydra vulgaris cells. Europhys. Lett. 64, 137– 143.

Golde, T., Pensalfini, M., Chahare, N., Roca-Cusachs, P., Wiche, G., Charras, G. T., Arroyo, M. and Trepat, X. (2026). Dynamics of supracellular keratin bundling and nuclear uncaging in stretched epithelia. Nat. Phys. 22, 1436–1445.

Groot, S.R.de and Mazur, P. (2013). Non-equilibrium thermodynamics. Newburyport: Dover Publications.

Grubor, N. M., Colovic, R. B., Atkinson, H. D. and Micev, M. T. (2013). Giant biliary mucinous cystadenoma of the liver. Ann Hepatol 12, 979–983.

Guo, Z., Yao, J., Zheng, X., Cao, J., Lv, X., Gao, Z., Guo, S., Li, H., Guan, D., Li, L., et al. (2025). Cavity oscillation drives pattern formation in early mammalian embryos. Cell Reports 44, 115342.

Ismail, N., Myint, K., Khaing, S. L., Giribabu, N. and Salleh, N. (2023). Cervical fluid pH, electrolytes and osmolarity changes and expression of ion transporters (ENaC, CFTR and AQP) in cervix of women with primary unexplained infertility. Mol Biol Rep 50, 6729–6737.

Kawagishi, R., Tahara, M., Sawada, K., Morishige, K., Sakata, M., Tasaka, K. and Murata, Y. (2004). Na+/H+ Exchanger-3 is involved in mouse blastocyst formation. Journal of Experimental Zoology 301A, 767–775.

Kennard, A. S., Sathe, M., Labuz, E. C., Prinz, C. K. and Theriot, J. A. (2023). Post-injury hydraulic fracturing drives fissure formation in the zebrafish basal epidermal cell layer. Current Biology 33, 2616-2631.e5.

Korotkevich, E., Niwayama, R., Courtois, A., Friese, S., Berger, N., Buchholz, F. and Hiiragi, T. (2017). The Apical Domain Is Required and Sufficient for the First Lineage Segregation in the Mouse Embryo. Dev Cell 40, 235-247.e7.

Latorre, E., Kale, S., Casares, L., Gómez-González, M., Uroz, M., Valon, L., Nair, R. V., Garreta, E., Montserrat, N., del Campo, A., et al. (2018). Active superelasticity in three-dimensional epithelia of controlled shape. Nature 563, 203– 208.

Lee, B. H., Fuji, K., Petzold, H., Seymour, P., Yennek, S., Schewin, C., Lewis, A., Riveline, D., Hiraiwa, T., Sano, M., et al. (2026). Permeability-driven pressure and cell proliferation control lumen morphogenesis in pancreatic organoids. Nat Cell Biol 28, 113–124.

Lee, B. H., Mukenhirn, M., Guyomar, T., Maggipinto, Y., Tanida, S., Fuji, K., Lu, L., Romer, F., Nonomura, M., Hiraiwa, T., et al. (2026). Biophysics of lumen morphogenesis. Development 153, dev205628.

Lenne, P.-F., Munro, E., Heemskerk, I., Warmflash, A., Bocanegra-Moreno, L., Kishi, K., Kicheva, A., Long, Y., Fruleux, A., Boudaoud, A., et al. (2021). Roadmap for the multiscale coupling of biochemical and mechanical signals during development. Phys. Biol. 18, 041501.

Leonavicius, K., Preece, C., Davies, B. and Srinivas, S. (2018). Mechanics of mouse blastocyst hatching revealed by a hydrogel-based microdeformation assay. Proceedings of the National Academy of Sciences of the United States of America 115, 10375–10380.

Manejwala, F. M., Cragoe, E. J. and Schultz, R. M. (1989). Blastocoel expansion in the preimplantation mouse embryo: Role of extracellular sodium and chloride and possible apical routes of their entry. Developmental Biology 133, 210–220.

Mazur, P. and Schneider, U. (1986). Osmotic responses of preimplantation mouse and bovine embryos and their cryobiological implications. Cell biophysics 8, 259–285.

Menezo, Y. and Guerin, P. (1997). The mammalian oviduct: biochemistry and physiology. European Journal of Obstetrics & Gynecology and Reproductive Biology 73, 99–104.

Mizobe, Y., Kuwatsuru, Y., Kuroki, Y., Fukumoto, Y., Tokudome, M., Moewaki, H., Aihoshi, S., Uenohori, K., Orita, Y., Iwakawa, T., et al. (2026). One-step warming of human blastocysts under different thawing solution volumes, processing times and osmotic pressures. Reproductive BioMedicine Online 52, 105430.

Mosaliganti, K. R., Swinburne, I. A., Chan, C. U., Obholzer, N. D., Green, A. A., Tanksale, S., Mahadevan, L. and Megason, S. G. (2019). Size control of the inner ear via hydraulic feedback. eLife 8, e39596.

Murayama, Y., Mizuno, J., Kamakura, H., Fueta, Y., Nakamura, H., Akaishi, K., Anzai, K., Watanabe, A., Inui, H. and Omata, S. (2006). Mouse zona pellucida dynamically changes its elasticity during oocyte maturation, fertilization and early embryo development. Human Cell 19, 119–125.

Muzumdar, M. D., Tasic, B., Miyamichi, K., Li, L. and Luo, L. (2007). A global double-fluorescent Cre reporter mouse. genesis 45, 593–605.

Niwayama, R., Moghe, P., Liu, Y.-J., Fabrèges, D., Buchholz, F., Piel, M. and Hiiragi, T. (2019). A Tug-of-War between Cell Shape and Polarity Controls Division Orientation to Ensure Robust Patterning in the Mouse Blastocyst. Developmental Cell 1–18.

Olver, D. J. and Benson, J. D. (2023). Meta-analysis of the Boyle van ‘t Hoff relation: Turgor and leak models explain non-ideal volume equilibrium. Cryobiology 113, 104581.

Onsager, L. (1931). Reciprocal Relations in Irreversible Processes. I. Phys. Rev. 37, 405–426.

Palmer, N. and Kaldis, P. (2016). Regulation of the Embryonic Cell Cycle During Mammalian Preimplantation Development. In Current Topics in Developmental Biology, pp. 1–53. Elsevier.

Pedersen, R. A. and Spindle, A. I. (1980). Role of the blastocoele microenvironment in early mouse embryo differentiation. Nature 284, 550–552.

Pinheiro, D., Hannezo, E., Herszterg, S., Bosveld, F., Gaugue, I., Balakireva, M., Wang, Z., Cristo, I., Rigaud, S. U., Markova, O., et al. (2017). Transmission of cytokinesis forces via E-cadherin dilution and actomyosin flows. Nature 545, 103–107.

Płusa, B. and Piliszek, A. (2020). Common principles of early mammalian embryo self-organisation. Development 147, dev183079.

Powers, R. D., Borland, R. M. and Biggers, J. D. (1977). Amiloride-sensitive rheogenic Na+ transport in rabbit blastocyst. Nature 270, 603–604.

Robinson, D. H., Bubien, J. K., Smith, P. R. and Benos, D. J. (1991). Epithelial sodium conductance in rabbit preimplantation trophectodermal cells. Dev Biol 147, 313–321.

Roffay, C., Molinard, G., Kim, K., Urbanska, M., Andrade, V., Barbarasa, V., Nowak, P., Mercier, V., García-Calvo, J., Matile, S., et al. (2021). Passive coupling of membrane tension and cell volume during active response of cells to osmosis. Proc Natl Acad Sci USA 118, e2103228118.

Rose, K. and Watson, A. J. (2007). Na/K-ATPase beta1 subunit expression is required for blastocyst formation and normal assembly of trophectoderm tight junction-associated proteins. The Journal of biological chemistry 282, 12127– 12134.

Rozema, D. and Maître, J.-L. (2024). Forces Shaping the Blastocyst. Cold Spring Harb Perspect Biol a041519.

Ruiz-Herrero, T., Alessandri, K., Gurchenkov, B. V., Nassoy, P. and Mahadevan, L. (2017). Organ size control via hydraulically gated oscillations. Development 144, 4422–4427.

Ryan, A. Q., Chan, C. J., Graner, F. and Hiiragi, T. (2019). Lumen Expansion Facilitates Epiblast-Primitive Endoderm Fate Specification during Mouse Blastocyst Formation. Developmental Cell.

Schliffka, M. F., Dumortier, J. G., Pelzer, D., Mukherjee, A. and Maître, J.-L. (2024). Inverse blebs operate as hydraulic pumps during mouse blastocyst formation. Nat Cell Biol.

Schliffka, M. F., Tortorelli, A.-F., Özgüç, Ö., de Plater, L., Polzer, O., Pelzer, D. and Maître, J.-L. (2021). Multiscale analysis of single and double maternal-zygotic Myh9 and Myh10 mutants during mouse preimplantation development. eLife 10, e68536.

Schindelin, J., Arganda-Carreras, I., Frise, E., Kaynig, V., Longair, M., Pietzsch, T., Preibisch, S., Rueden, C., Saalfeld, S. and Schmid, B. (2012). Fiji: an open-source platform for biological-image analysis. Nature Methods 9, 676–682.

Shah, A. A., Soomro, N. A., Talib, R. K., Sadhayo, A. N. and Soomro, S. A. (2014). Giant intraabdominal endometrial cyst. J Coll Physicians Surg Pak 24, 438–440.

Shi, W., Duclut, C., Xu, Y., Ma, Y., Qiao, J., Lin, B., Yang, D., Prost, J. and Dong, B. (2025). Physics of notochord tube expansion in ascidians. Proc. Natl. Acad. Sci. U.S.A. 122, e2419960122.

Sofroniew, N., Lambert, T., Bokota, G., Nunez-Iglesias, J., Sobolewski, P., Sweet, A., Gaifas, L., Evans, K., Burt, A., Doncila Pop, D., et al. (2026). napari: a multi-dimensional image viewer for Python.

Stokkermans, A., Chakrabarti, A., Subramanian, K., Wang, L., Yin, S., Moghe, P., Steenbergen, P., Mönke, G., Hiiragi, T., Prevedel, R., et al. (2022). Muscular hydraulics drive larva-polyp morphogenesis. Current Biology 32, 4707-4718.e8.

Swinburne, I. A., Mosaliganti, K. R., Upadhyayula, S., Liu, T.-L., Hildebrand, D. G. C., Tsai, T. Y.-C., Chen, A., Al-Obeidi, E., Fass, A. K., Malhotra, S., et al. (2018). Lamellar projections in the endolymphatic sac act as a relief valve to regulate inner ear pressure. eLife 7,.

Timbs, M. M. and Spring, K. R. (1996). Hydraulic Properties of MDCK Cell Epithelium. Journal of Membrane Biology 153, 1–11.

Torres-Sánchez, A., Kerr Winter, M. and Salbreux, G. (2021). Tissue hydraulics: Physics of lumen formation and interaction. Cells & Development 168, 203724.

van’t Hoff, J. H. (1884). Etudes de dynamique chimique. Amsterdam : Frederik Muller.

Venkova, L., Vishen, A. S., Lembo, S., Srivastava, N., Duchamp, B., Ruppel, A., Williart, A., Vassilopoulos, S., Deslys, A., Garcia Arcos, J. M., et al. (2022). A mechano-osmotic feedback couples cell volume to the rate of cell deformation. eLife 11, e72381.

Wassarman, P. M. and Mortillo, S. (1991). Structure of the Mouse Egg Extracellular Coat, the Zona Pellucida. In International Review of Cytology, pp. 85–110. Elsevier.

Weevers, S. L., Falconer, A. D., Mercker, M., Sadeghi, H., Rozema, D., Ferenc, J., Maître, J.-L., Ott, A., Oelz, D. B., Marciniak-Czochra, A., et al. (2025). Mechanochemical patterning localizes the organizer of a luminal epithelium. Sci. Adv. 11, eadu2286.

Wiley, L. M. (1984). Cavitation in the mouse preimplantation embryo: Na/K-ATPase and the origin of nascent blastocoele fluid. Dev Biol 105, 330–342.

Wu, H., Duclut, C., Arkowitz, G., Chilupuri, R., Dang, T., Prost, J., Ladoux, B. and Mège, R.-M. (2025). Regulation of epithelial tissue homeostasis by active transepithelial transport. Proc. Natl. Acad. Sci. U.S.A. 122, e2503156122.

Yang, Q., Xue, S.-L., Chan, C. J., Rempfler, M., Vischi, D., Maurer-Gutierrez, F., Hiiragi, T., Hannezo, E. and Liberali, P. (2021). Cell fate coordinates mechano-osmotic forces in intestinal crypt formation. Nat Cell Biol 23, 733–744.

Yoshiba, S., Shiratori, H., Kuo, I. Y., Kawasumi, A., Shinohara, K., Nonaka, S., Asai, Y., Sasaki, G., Belo, J. A., Sasaki, H., et al. (2012). Cilia at the Node of Mouse Embryos Sense Fluid Flow for Left-Right Determination via Pkd2. Science (New York, NY).

Yu Sun, Kai-Tak Wan, Roberts, K. P., Bischof, J. C. and Nelson, B. J. (2003). Mechanical property characterization of mouse zona pellucida. IEEE Trans.on Nanobioscience 2, 279– 286.

Zenker, J., White, M. D., Gasnier, M., Álvarez, Y. D., Lim, H. Y. G., Bissiere, S., Biro, M. and Plachta, N. (2018). Expanding Actin Rings Zipper the Mouse Embryo for Blastocyst Formation. Cell 1–34.

## SUPPLEMENTARY REFERENCES

[1] L. Onsager, Reciprocal Relations in Irreversible Processes. I., Phys. Rev. 37, 405 (1931).

[2] S. De Groot and P. Mazur, Non-Equilibrium Thermodynamics (Dover Publications, New York, U.S.A., 1984).

[3] A. Torres-Sánchez, M. Kerr Winter, and G. Salbreux, Tissue hydraulics: Physics of lumen formation and interaction, Cells & Development 168, 203724 (2021).

[4] K. Leonavicius, C. Royer, C. Preece, B. Davies, J. S. Biggins, and S. Srinivas, Mechanics of mouse blastocyst hatching revealed by a hydrogel-based microde formation assay, Proc. Natl. Acad. Sci. U.S.A. 115, 10375 (2018).

[5] J. G. Dumortier, M. Le Verge-Serandour, A. F. Tortorelli, A. Mielke, L. de Plater, H. Turlier, and J.-L. Maître, Hydraulic fracturing and active coarsening position the lumen of the mouse blastocyst, Science 365, 465 (2019).

[6] C. J. Chan, M. Costanzo, T. Ruiz-Herrero, G. Mönke, R. J. Petrie, M. Bergert, A. Diz-Muñoz, L. Mahadevan, and T. Hiiragi, Hydraulic control of mammalian embryo size and cell fate, Nature 571, 112 (2019).

